# Millimeter-scale selective amplification in the developing visual cortex

**DOI:** 10.64898/2026.03.31.715687

**Authors:** Haleigh N. Mulholland, Sigrid Trägenap, Matthias Kaschube, Gordon B. Smith

## Abstract

Cortical processing is believed to involve the selective amplification of specific inputs by recurrent networks. But whether selective amplification operates within the millimeter-scale networks underlying perception and behavior remains unclear. Here, we combine patterned optogenetic stimulation—informed by computational models—with calcium imaging in immature ferret visual cortex to show that cortical networks are preferentially activated by inputs aligned to endogenous recurrent subnetworks. Dominant subnetworks can be inferred from spontaneous activity, where response reliability and specificity depend on the overlap of input with these dominant modes. Well-aligned inputs are selectively amplified, evoking more reliable and specific responses that exhibit greater spatiotemporal stability than misaligned inputs. This preferential amplification suggests early cortical networks leverage endogenous dynamics to stabilize and refine precise sensory representations over development.

## Introduction

Sensory perception and behavior recruit neural activity distributed over large-scale networks that can span millimeters of the cerebral cortex. These networks are made up of dense and highly recurrent connections that can profoundly shape responses to cortical inputs (*1–5*). Thus, in order to understand how such mesoscopic functional networks operate, it is crucial to determine how incoming patterns of activity interact with these recurrent cortical circuits. Theoretical work indicates that certain inputs can engage with recurrent connections more effectively than others, resulting in selective amplification of activity within preferentially coupled subnetworks (*5–9*). Within such models, selective amplification plays a critical role in shaping response selectivity and reliability(*6–8, 10, 11*), and has important implications for learning (*12, 13*), allowing for the formation and maintenance of robust sensory and behavioral representations.

Previously, cortical interactions within recurrent networks have been investigated on a local scale in mature cortical circuits, using cellular resolution imaging and targeted stimulation of subpopulations within several hundred microns, revealing selectively coupled local ensembles that recurrently modulate activity (*3, 5, 14, 15*). However, currently little is known about the principles determining whether specific input patterns effectively engage amplification in the columnar functional networks that are a hallmark of cortical organization in humans, non-human primates, and carnivores (*16–19*). In these networks—such as the orientation preference maps found in V1— cortical columns made up of neighboring neurons with similar response properties are classically considered to be the ‘functional units’ of neural processing(*18, 19*), which together form mesoscopic, distributed networks spanning several millimeters of cortex. Selective amplification in millimeter-scale networks is supported by computational models predicting selective biases for inputs aligned to recurrent circuitry, as compared to random and misaligned inputs (Fig. 1A) (*20, 21*), but directly testing this *in vivo* has been technically challenging. Therefore, we currently do not have a thorough understanding of the nature of network amplification at this scale of cortical processing, nor do we know what patterns are most effective in driving selective amplification in mesoscale cortical networks.

**Figure 1.**
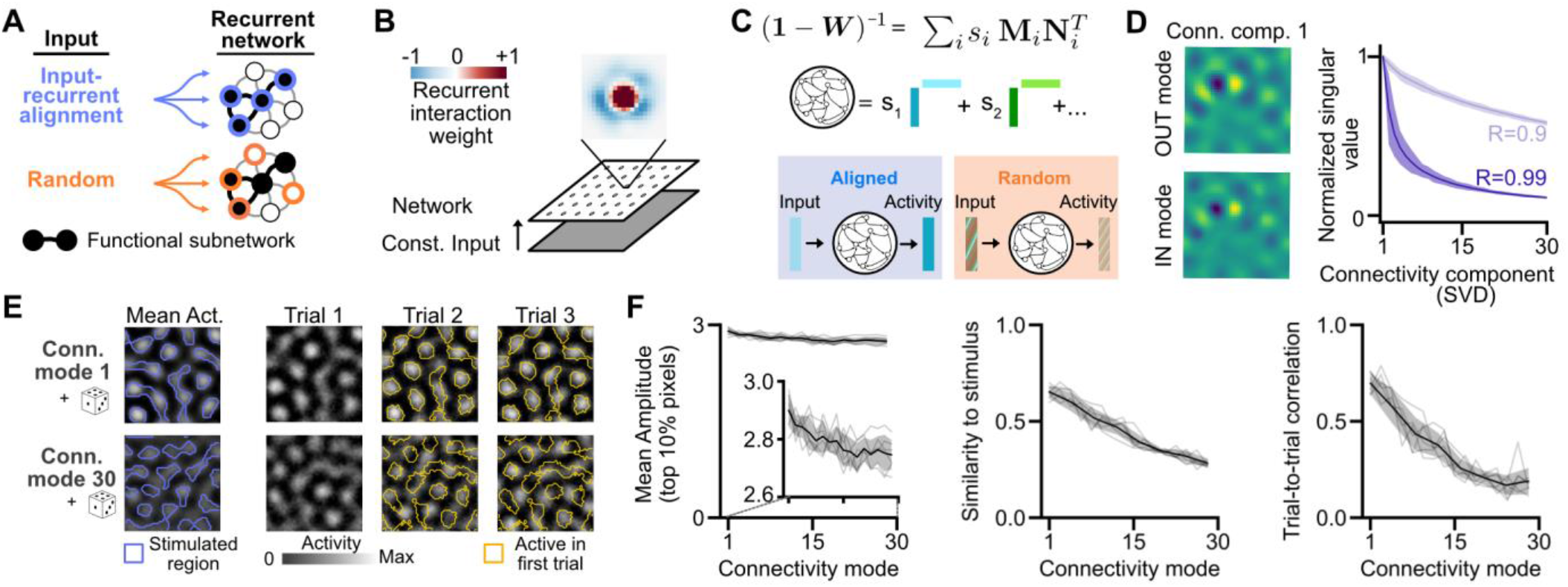
Stimulus patterns close to dominant connectivity modes drive reliable network responses *in silico*. **(A)** Schematic: Inputs into a recurrent network can either align (input-recurrent alignment, *top*) with functional subnetworks or be randomly targeted (*bottom*). **(B)** Modeling schematic: Recurrent network with lateral excitation and long-range inhibition (LE/LI) interaction profile with a mild degree of additional heterogeneity. **(C)** Schematic: Decomposition of recurrent network interactions into input- and output modes based on Singular Value Decomposition. **(D)** Example IN- and OUT-pattern of the first connectivity mode (*left*) with associated singular values for the first thirty modes for two different recurrent strengths (*right*; strong: R=0.99, moderately weaker: R=0.9). Thick line: average over N=50 networks, shaded region: standard deviation. **(E)** (*Left*) Driving network with patterns based on connectivity modes and additional noise. (*Middle*) Mean evoked network response across trials (n=40 trials), contours mark stimulated region. (*Right*) Example network activity patterns, contours mark active region in first trial. **(F)** Stimulus patterns based on leading connectivity modes (high associated singular values) evoke responses with greater peak amplitude (*left*), greater similarity to stimulus (*middle*), and higher trial-to-trial correlations (*right*) compared to higher order connectivity modes with weaker associated singular values. Individual lines: N=6 independent networks. Thick line and shaded region show mean and standard deviation.

Here we address this question through targeted optogenetic stimulation of cortical networks *in vivo* in developing ferret visual cortex. We take advantage of the human-like, modular (or ‘columnar’) functional networks that exist in this species in both sensory and non-sensory areas (*17, 22, 23*), which allows us to use single-photon optogenetic stimulation to selectively drive networks with specific patterned inputs over several millimeters (*24, 25*). Importantly, activity patterns in the developing ferret cortex appear to be dominated by recurrent interactions (*20, 25, 26*), as opposed to feedforward, providing an ideal system to study the effect of local circuits on network amplification.

Using a combination of computational modeling and *in vivo* experiments, we show that amplification by the millimeter-scale networks in visual cortex is highly input-selective and primarily affects response reliability rather than magnitude. Model simulations predict that the inputs most susceptible to amplification correspond to the dominant modes of connectivity of the network and that these modes can be effectively estimated from the dominant modes of spontaneous activity, thereby allowing for tractable experiments *in vivo*. By applying patterned optogenetic stimuli *in vivo*, we find that inputs well-aligned with spontaneous activity evoke responses that are more reliable, specific, and show more stable spatiotemporal dynamics than random stimuli, with the strength of these effects predicted by the degree of alignment. Our results demonstrate millimeter-scale selective amplification and suggest that this is driven via local recurrent interactions. Such a mechanism in early development suggests a potential means by which developing cortical networks can build the selective and reliable responses seen in the mature brain.

## Results

### Model predicts most reliable responses for inputs aligned with leading connectivity modes

To investigate the impact of recurrent amplification in developing millimeter-scale cortical networks, we began with a computational model. Expanding on previous work(*20, 25, 26*), we studied a dynamic, non-linear, firing-rate network, implementing random recurrent connectivity with a local-excitation/lateral-inhibition (LE/LI) structure and a moderate degree of spatial heterogeneity (Fig. 1B), which has been shown to recapitulate the large-scale and modular spontaneous activity patterns seen in developing ferret cortex (*26*).

Hypothesizing that the best amplified inputs could be estimated from the connectivity structure of the network (Fig. 1A), we sought to examine this connectivity by performing singular value decomposition (SVD) of the linearized network under static input (Fig. 1C). This analysis decomposes the effective recurrent interactions into independent, rank one connectivity components, each one composed of a pair of IN/OUT connectivity modes and a singular value whose magnitude expresses the relative contribution of each component (Fig. 1C, and Methods). A stimulus aligned with the IN-connectivity mode associated with the largest singular value is thus predicted to result in maximal possible amplification in a linear network, and the resulting activity is proportional to the OUT-connectivity mode. In our spatial network, both modes can be displayed as spatial patterns (Fig. 1D, left). Stimuli overlapping with the first few leading (IN) connectivity modes are expected to show greater amplification than random patterns or patterns overlapping higher-order connectivity modes. Moreover, if the network is nearly symmetric—i.e. interactions are primarily reciprocal— corresponding IN- and OUT-connectivity modes are similar, and thus driving such a mode results in an activity pattern similar to the input drive (Fig. 1C, lower left). However, if not symmetric, or if a stimulus overlaps with many connectivity modes that vary greatly in their singular values (e.g. a random input, Fig. 1C lower right), the resulting output can differ strongly from the stimulus pattern.

We found that in our model network, owing to its LE/LI recurrent structure, the leading connectivity modes are modular and distributed across space (Fig. 1D). Moreover, despite this common modular structure, the leading connectivity modes vary considerably in their singular values (Fig. 1D, right). Thus, unlike in the case of spatially homogeneous recurrent connectivity (Fig. S1A,B), there is a lack of degeneracy of connectivity modes in our model, predicting selective amplification in millimeter-scale cortical networks is highly dependent on the specific spatial structure of the stimulus pattern. Finally, IN- and OUT-connectivity modes are notably similar in our model (Fig. 1D, left), indicating that, despite heterogeneity in its recurrent connections, the network’s most amplifiable patterns are shaped through nearly symmetric interaction motifs. The model thus predicts that an external stimulus applied to the millimeter-scale cortical network produces an output activity pattern that closely resembles the input pattern provided that the input is aligned to the dominant connectivity modes in the network.

To test whether these properties hold also in the full nonlinear model, we stimulated with inputs based on the SVD-derived connectivity modes of the network, while adding a noise component to the input to model trial-to-trial variability (see Methods). We found that stimulation patterns based on connectivity modes evoke robust modular responses that closely match the input patterns when averaged across trials (Fig. 1E, left). For the leading connectivity mode, evoked responses are spatially similar to their input and are remarkably consistent across trials (Fig. 1E, top). However, higher order connectivity modes evoke outputs with greater trial-to-trial variation, suggesting that these inputs weakly engage the network and are susceptible to noise (Fig. 1E, bottom).

A systematic analysis revealed that whereas the amplitude of individual responses decreases only slightly for higher order connectivity modes (Fig. 1F left), the similarity of single trial responses to the input pattern (Fig. 1F middle) and, in particular, the average trial-to-trial correlation between these responses (Fig. 1F right) depend sensitively on the order of the connectivity mode. Thus, inputs that align with dominant connectivity modes revealed by SVD preferentially engage the large-scale recurrent network in the full nonlinear model. However, rather than producing strong changes in the amplitude of responses, as expected in a linear model (*27–29*) (see Fig. S2), this engagement manifests most clearly in increased response reliability and similarity to input (see Fig. S3 for robustness against parameter variations). Furthermore, our results suggest that these response properties could provide a sensitive readout of selective network amplification that we can now examine *in vivo*.

### Inputs derived from endogenous activity produce reliable responses

While these results indicate that preferred inputs can be derived from network connectivity, obtaining detailed connectivity at large scale *in vivo* is highly challenging. We therefore utilized our network model to examine the possibility that dominant connectivity modes can be instead inferred from ongoing network activity. We reasoned that if the modular, low-dimensional structure of spontaneous activity in ferret visual cortex emerges from local intracortical interactions, as indicated by recent experiments (*25*), then spontaneous activity should carry signatures of the dominant modes of recurrent connectivity. Thus, spontaneous correlation maps—which reveal which modules are typically co-active—could serve as an experimentally tractable proxy for the network’s dominant connectivity modes.

To explore this idea in the model, we assumed spontaneous activity arises from broadly distributed inputs, which has been shown to account for several key features of spontaneous activity in developing ferret visual cortex (*20, 25, 26*). We therefore computed spontaneous correlation patterns ((*26*), see Methods) from model simulations (Fig. 2A)—referred to as ‘endogenous’ patterns in the following—and found that these indeed tend to overlap more strongly with the leading connectivity modes derived from the SVD of the recurrent network than control patterns consisting of random bandpass filtered noise with matched spatial statistics (Fig. 2B, left). To quantify this alignment to connectivity, we averaged over the correlation with individual IN-connectivity modes weighted by their singular value (see Methods), finding that endogenous patterns are significantly more aligned than the random control patterns (Fig. 2B, right). When examining the activity patterns evoked by these stimuli in our model, we found that endogenous patterns evoke activity that is slightly stronger in amplitude and much more reliable and similar to their inputs as compared to the random control inputs (Fig. 2C-E), suggesting that inputs based on spontaneous activity can indeed be used to effectively engage dominant modes of connectivity.

**Figure 2.**
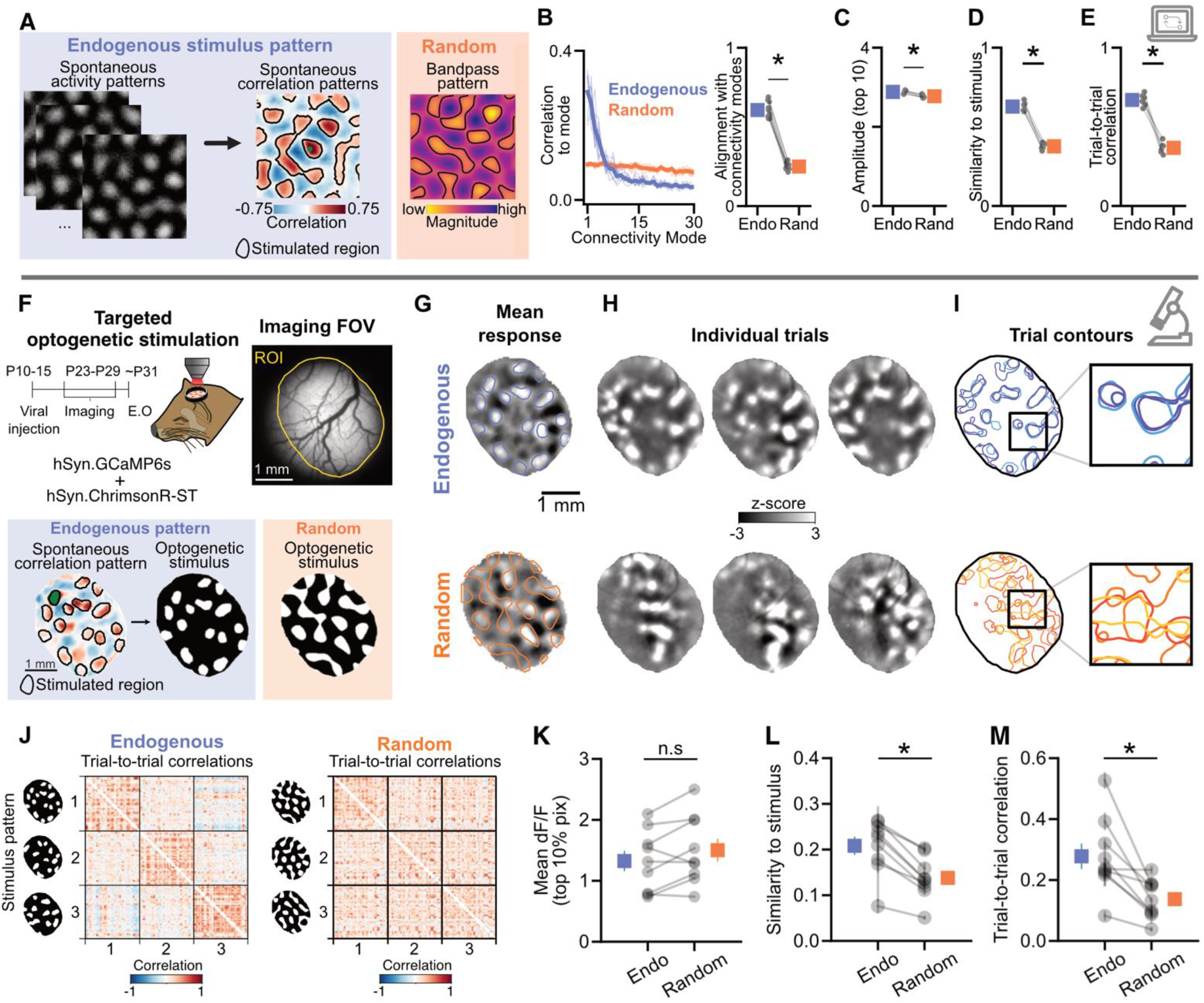
Selective amplification of stimulus patterns based on endogenous modes of activity drives more reliable cortical responses than random patterns. **(A)** Schematic of model network inputs based on endogenous spontaneous correlation patterns (derived from spontaneous activity patterns driven by uniform noise) (*left*) and random bandpass filtered noise (*right*). **(B)** Endogenous-derived patterns show more overlap (correlation between stimulus pattern and i-th connectivity mode) with dominant connectivity modes than do random patterns (*left*). Thin lines indicate n=10 individual networks, averaged over n=200 stimuli for endogenous derived (blue) and random (orange). Thick lines show average. Alignment with connectivity modes (*right*) quantifies this sum of the overlap weighted with the associated singular value, where grey dots are n=10 independent networks, and error bars indicate mean +/− std. Legend also applies to C-E. (**C-E**) Stimulus patterns based on spontaneous correlation patterns evoke responses with greater amplitude (**C**) and similarity to stimulus (**D**), and trial-to-trial correlation (**E**), compared to random stimuli (n=6 independent simulations). **(F)** *In vivo* experimental schematic: wide-field targeted optogenetics and calcium imaging in postnatal ferret visual cortex. Optogenetic stimulus patterns were designed based on spontaneous correlation patterns (*left*, blue, endogenous) or random bandpass-filtered noise (*right*, orange, random). **(G)** Mean response to an example endogenous (*top*, blue) and random (*bottom*, orange) input pattern. Contours of stimulus pattern overlaid to show stimulus-specific responses (n=40 trials per pattern). **(H)** Example individual trial responses, showing top three trials with greatest similarity to stimulus. **(I)** Contours of individual trials in D. Endogenous-based inputs evoke highly similar patterns across trials, while random inputs show greater spatial variability. **(J)** Evoked activity trial-to-trial correlation matrix for 3 different endogenous (*top*) and random (*bottom*) stimulus patterns in an example animal. **(K)** Average ΔF/F response within most active pixels (top 10%) on a per trial basis for endogenous- and random-evoked responses (n=9 animals, p=0.07, WSR test). Grey circles: mean across input patterns (n=3-4) for each stimulus type for individual animals. Colored squares: Group average. Error bars=+/− SEM). Legend also applies to L, M. **(L)** Average correlation of individual trials to their respective stimulus patterns (n=9 animals, p<0.01, WSR test). **(M)** Trial-to-trial correlation, averaged for each stimulus pattern (n=9 animals, p<0.01, WSR test).

Guided by our model’s prediction that developing cortical networks are preferentially driven by input patterns matching those present in correlated patterns of spontaneous activity, we next sought to test this *in vivo* through direct control of input activity in layer 2/3 networks via patterned optogenetic stimulation. Notably, the modular nature of the dominant connectivity modes in computational models that recapitulate activity in the developing cortex (Fig. 1D), together with the columnar structure of correlated spontaneous activity *in vivo* allows us to use 1-photon millimeter-scale patterned optogenetic stimulation to test this prediction. Thus, we performed widefield calcium imaging and patterned single photon optogenetics in young ferret visual cortex prior to eye opening in animals expressing GCaMP6s and ChrimsonR-ST in excitatory neural populations. Stimulation was delivered using our custom-designed opto-macroscope (*24*), which allowed us to project patterned input activity and measure responses over approximately a 3 mm diameter field-of-view (Fig. 2F, left). To design input patterns putatively aligned to dominant network connectivity modes, we first imaged spontaneous activity and then identified frequently occurring spontaneous patterns via online analysis of spontaneous correlation maps (Fig. 2F, right; see Methods). We then created optogenetic stimulation patterns based on these maps (Fig. 2F, bottom left, ‘endogenous’, blue, Fig. S4). As a control, we also generated random patterns with low expected alignment to the endogenous network by bandpass filtering white noise at a wavelength corresponding to the characteristic spatial wavelength of activity observed in V1 activity (*22, 25, 30*) (Fig. 2F, bottom right, ‘random’, orange). To test the specificity of evoked responses, multiple (minimum 3) different stimulus patterns of each category were interleaved within an imaging session.

We observed that both endogenous and random stimuli drove robust modular patterns of activity that in trial-averaged responses aligned well with the input stimulus (Fig. 2G, S4, S5). However, when examining the individual trials these two types of input showed marked differences in the reliability of the evoked pattern (Fig. 2H-J). As predicted by our model, the strength of individual trial evoked responses was similar regardless of input type (Fig. 2K; SI2; endogenous=1.32 ± 0.17, random=1.50 ± 0.19, trial averaged ΔF/F ± SEM within top 10% most active pixels on each trial, n=3-4 input patterns across n=9 animals, p=0.07, WSR test, Fig. S6), but endogenous inputs evoked activity patterns that tended to be more consistent trial-to-trial than the random control patterns (Fig. 2M; S5; endogenous=0.28 ± 0.04, random=0.14 ± 0.02, mean ± SEM, p<0.01, WSR test). Endogenous-based stimuli also drove responses that were more similar to their inputs than random stimuli (Fig. 2L; S5; endogenous=0.21 ± 0.02, random=0.14 ± 0.02, p<0.01, WSR test).

Notably, this difference in amplification between endogenous and random stimuli was substantial compared to the effect we observed when varying the spatial wavelength of random input patterns. In networks with LE/LI interactions, such a dependency on the input’s spatial wavelength is expected due to the tendency of LE/LI interactions to promote activity around a characteristic spatial wavelength (*25*). We confirmed such relationship in our full nonlinear model (Fig. S7A-C, orange) and observed a consistent trend *in vivo* for random stimuli of different wavelengths (Fig. S7 D-F, orange). However, even when the stimulus wavelength matched the network’s intrinsic scale evidenced by the characteristic wavelength of spontaneous activity (Fig. S7 D), endogenous inputs still evoked considerably more reliable and input-aligned responses than random controls (Fig. S7 B,C,E,F, blue). Thus, while matching the network’s intrinsic spatial wavelength is a prerequisite for the effective amplification of an input, it is insufficient to account for the high reliability we observed when driving the cortex with endogenous input patterns. Rather, our modeling results show that even a moderate degree of symmetry-breaking heterogeneity within LE/LI recurrent interactions—any deviation from perfect isotropy— results in selective amplification beyond that driven by wavelength amplification alone (compare Fig. 1F with Fig. S1D), consistent with the notion that such heterogeneity creates competition between preferentially coupled subnetworks (*26*).

Together, these empirical results corroborate our model predictions and show that developing networks exhibit preferential biases towards inputs similar to endogenous activity over random ones, with the former driving more specific and reliable responses, thus demonstrating the presence of selective amplification at the millimeter scale in the visual cortex.

### Endogenous based inputs evoke temporally stable responses

One potential advantage of systems exhibiting selective amplification is that outputs may be less susceptible to incidental temporal noise and thus exhibit greater stability over time in response to sustained inputs. To test whether this is the case in our data and to determine how this stable recurrent amplification is influenced by the alignment of inputs with the dominant modes of the network, we added time-varying, temporally correlated noise to the inputs in our computational model (Fig. 3A). We found that sustained drive with endogenous input patterns elicit activity that is more stable (Fig. 3B-C) and deviates less from its inputs over time than that evoked by random patterns (Fig. 3D). These model results predict that misaligned stimuli will drive modes that are more susceptible to the continuous influence of noise over time and therefore evoke a changing array of activity patterns during sustained stimulation.

**Figure 3.**
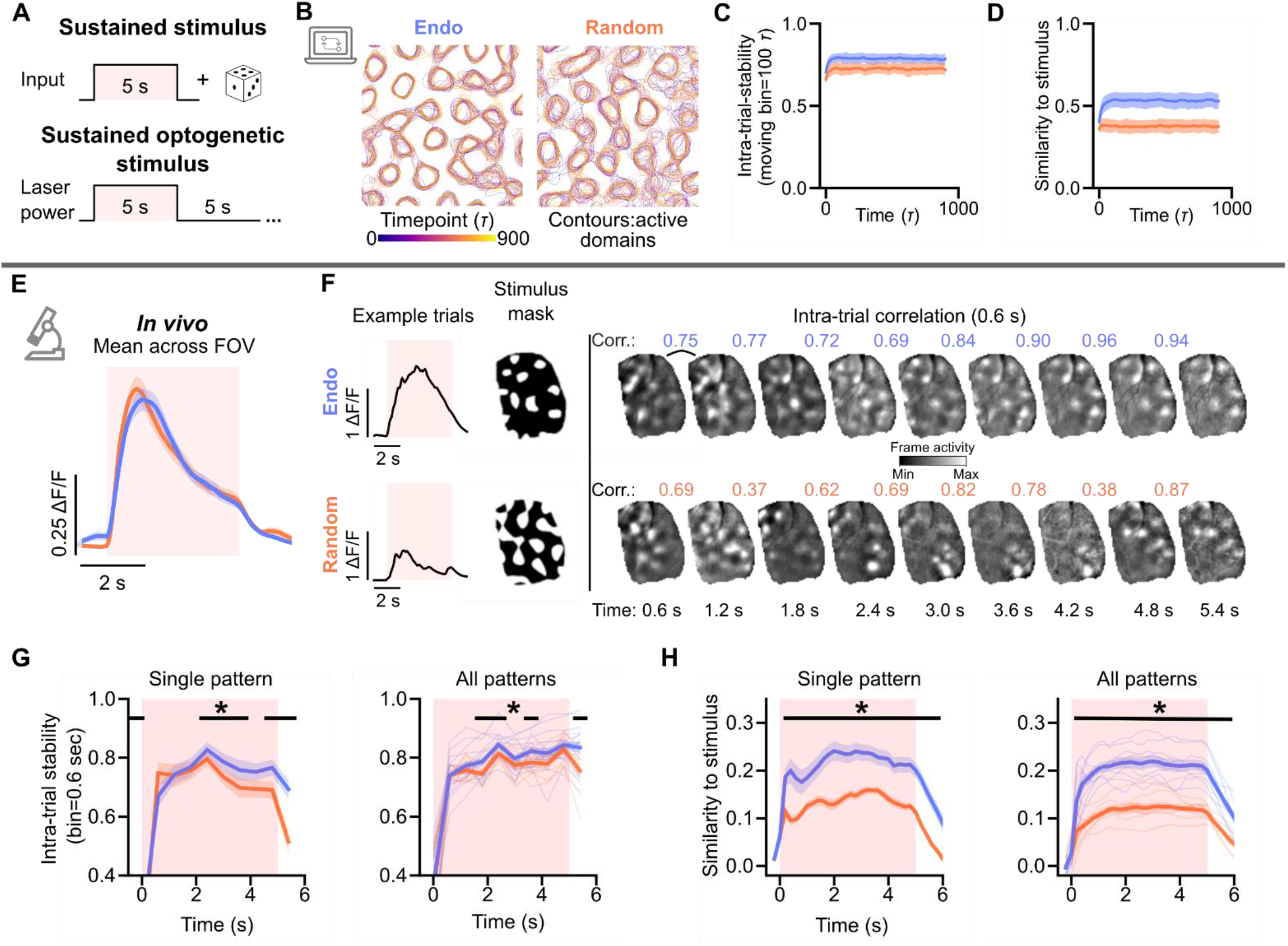
Endogenous-based stimuli evoke more stable responses to sustained input. **(A)** Schematic: (*Top*) Model simulation of sustained input using a fixed input drive and a time varying noise component. (*Bottom*) *In vivo* experimental design, with 5 seconds of continuous patterned optogenetic stimulation. **(B)** Evolution of activity patterns during 900 time steps (*τ*) for endogenous-derived stimulus (*left*) and random bandpass pattern (*right*). Contours show active domains during different time points (purple to yellow) in the simulation period. (**C-D**) Model simulation: Endogenous patterns drive responses with greater intra-trial stability (C) and are more similar to their stimulus (D) over long durations than random patterns. Thick lines and shaded region show mean and standard deviation across n=3 independent networks. **(E)** *In vivo* experiment: Stimulus triggered average ΔF/F across field of view (n=40 trials) for example endogenous and random stimuli. **(F)** Example evoked ΔF/F time series for an individual trial for two example stimulus patterns, endogenous (*top*) and random (*bottom*). (*Left*) Average ΔF/F across field of view evoked by given stimulus pattern. (*Right*) Individual, spatially filtered frames, showing the evolution of patterns over the duration of the stimulus. Frame-to-frame correlations for the example frames shown above. **(G)** Intra-trial stability of evoked activity, measured as the correlation between frames separated by 0.6 seconds. (*Left*) Mean (+/− SEM) single pattern stability across n=40 trials for examples shown in (E-F). Black bar indicates periods where p<0.05. (*Right*) Mean stability across all patterns (n=21 endogenous, 9 random patterns, pooled across n=4 animals). **(H)** Same as F, except for the correlation between evoked response and the stimulus input pattern.

To directly test this prediction *in vivo*, we optogenetically stimulated with endogenous and random patterns for a prolonged duration (5 seconds) in a subset of experiments. These sustained stimulations evoked responses with transient peaks approximately 1 second after stimulus onset but continued to produce elevated patterns of modular activity for the duration of the stimulus (Fig. 3E-F). On individual trials, we frequently observed multiple transient peaks in the activity throughout the stimulation period (Fig. 3F, left). We quantified the intra-trial stability of the evoked activity by computing frame-to-frame correlations over the course of the stimulation (Fig. 3F, right; approx. 0.6 second bins). Endogenous-based stimuli evoked patterns of activity that were significantly more stable over time relative to responses from random stimuli, which tended to evolve dynamically into different patterns over the duration of stimulation despite the unchanging input (Fig. 3F,G).

Additionally, the activity patterns evoked by endogenous stimuli were more similar to their stimulus inputs over time than those evoked by random inputs (Fig. 3H). Importantly, because both stimulus types elicited similarly strong activity (Fig. 3E), stimulus-specific variations in the calcium indicator’s decay kinetics are highly unlikely, and thus cannot account for the observed differences in response stability. Thus, these results demonstrate a stimulus-dependent effect on the temporal dynamics of network activity in the visual cortex, with inputs corresponding to endogenous activity driving more stable responses.

### Selectivity of network amplification predicted by similarity to spontaneous activity

In our modelling results, the strength of network amplification decreases in a graded fashion as a function of connectivity mode, with amplification strongest for inputs aligned to the leading modes (Fig. 1F). This selectivity is rooted in the network’s heterogeneity, as such a systematic trend is absent for isotropic, spatially homogenous connectivity (Fig. S1D). If this principle holds *in vivo*, we expect to observe that not all endogenous-derived patterns engage cortical networks equally. Rather, the degree to which any input pattern (endogenous or random) is amplified by the recurrent network should be determined by how well this input aligns with the underlying network connectivity. Moreover, if, as suggested by our modelling, spontaneous activity can be used as a proxy for a network’s connectivity modes, we expect that the degree of amplification can be predicted based on similarity to the leading principal components (PCs) of spontaneous activity, which explain the largest fraction of the variance and thus should sensitively reflect the underlying recurrent circuitry.

To test this idea, we first examined the selectivity of network amplification in our computational model. We found that stimulating the network with leading PC patterns derived from simulated spontaneous activity (see Fig. 2A and Methods) evokes responses that are highly reliable across trials and similar to the input, whereas those based on higher-order PCs evoke unreliable responses that are less similar to their input pattern (Fig 4A, B). This indicates that the network is biased towards amplifying input patterns that align with the dominant modes (PCs) of spontaneous activity, mirroring the dependency on connectivity modes observed earlier (Fig. 1F). Critically, when analyzing the random and endogenous inputs from Figs 2 and 3, we observed a tight relationship between their alignment to connectivity (computed as in Fig. 2B) and their alignment to spontaneous activity (determined analogously by the average correlation with all spontaneous activity PCs weighted by their explained variance; see Methods) (Fig. 4C), indicating that the latter is indeed an accurate proxy of the former. Using all input types—including endogenous and random, as well as inputs specifically constructed with varying alignment to spontaneous activity—our model simulations then show that alignment to spontaneous activity predicts the reliability of trial-to-trial correlations and similarity to stimulus, as well as the spatiotemporal stability of activity to sustained inputs (Fig 4 D-G).

**Figure 4.**
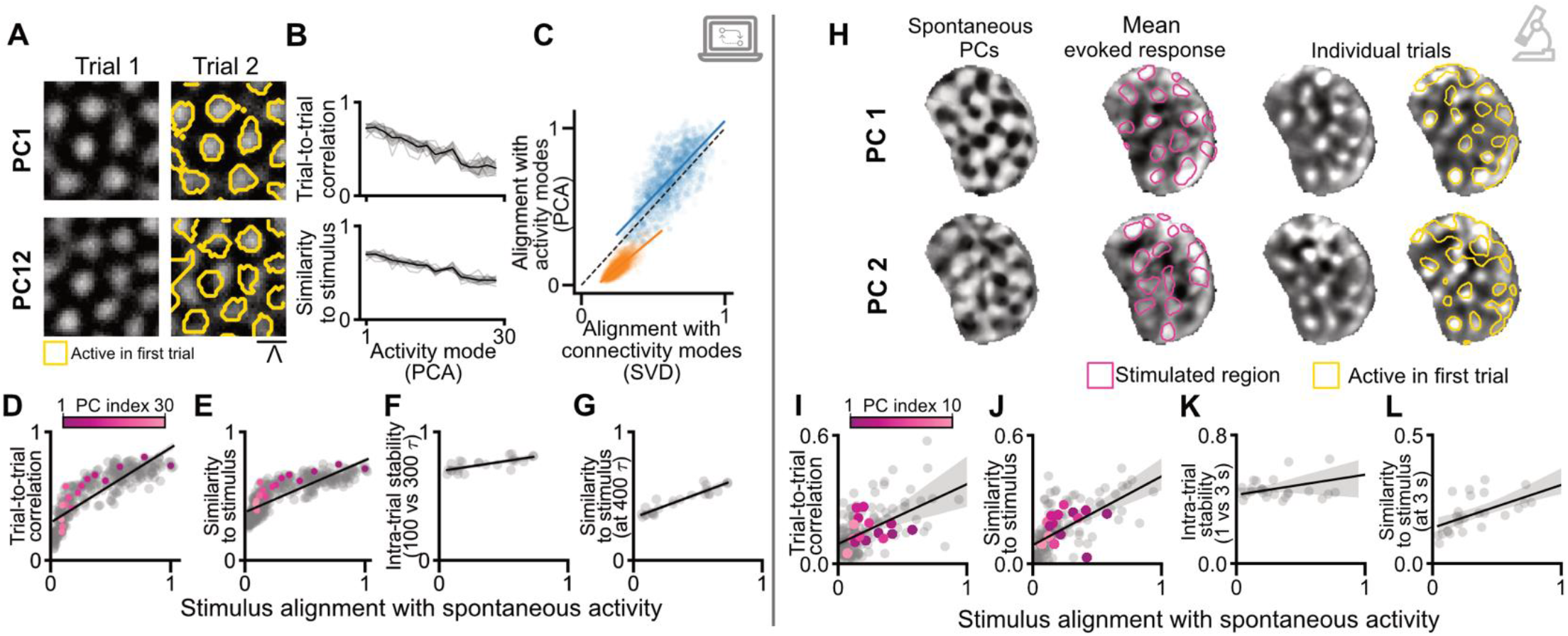
Degree of stimulus alignment with spontaneous principal components predicts selectivity of amplification. (**A-B**) Computational model of activity driven by inputs based on the principal components of spontaneous activity (derived from uniform input into network, see Methods). (**A**) Individual trial responses to leading and higher-order PCs. Leading PC patterns produce reliable trial-to-trial activity, while higher-order inputs have greater variability in spatial pattern. (Yellow: Contours of trial 1 active pixels). **(B)** Trial-to-trial correlation (*top*) and similarity to stimulus (*bottom*) as a function of inputs corresponding to increasing spontaneous PC index (n=6 simulations, mean, +/− SEM). **(C)** Alignment with network connectivity (SVD) modes (see figure 2B) compared to alignment with spontaneous activity (PCA) modes (see Methods) for endogenous (blue) and random stimuli (orange). Lines and shaded regions indicate linear regression best fit +/− 95% confidence interval. **(D)** Trial-to-trial correlation vs stimulus pattern alignment with spontaneous PCs. Each dot corresponds to the average across trials for a given stimulus pattern. Colored dots show example PC-based input patterns, hued by PC index (A). Black lines and shaded region indicate linear regression best fit +/− 95% confidence interval. **(E)** As (D), but for similarity to stimulus input. (**F**). As (D), but for intra-trial-stability between 100 and 300 *τ*. **(G)** As (D), but for similarity to stimulus at 400 *τ*. **(H)** *In vivo* data: For optogenetic stimulus patterns based on the principal components of spontaneous activity (*left*), mean response (*middle*) and example individual trials (*right*). Two example patterns based on PCs 1 and 2 shown, contours of inputs in magenta. (**I-L**) Same as D-G, but for *in vivo* data. (I-J) n=169 stimuli, pooled across 9 animals. (K-L) n=26 stimuli, pooled across 4 animals. Intra-trial-stability (K) was determined between 1 and 3 secs, similarity to stimulus (L) at 3 sec.

We then tested if such a unifying metric of alignment to spontaneous activity could explain the variability we observed *in vivo* across individual stimulus patterns and animals (Fig 2L-M) and predict why certain input patterns produce more reliable responses over others. To examine this, we pooled together stimulus patterns from all experiments (including endogenous and random stimuli of all wavelengths, Figs 2 and SI4) and calculated their alignment with spontaneous activity, which we computed, as for the model, based on the PCs of spontaneous activity. Additionally, for a subset of experiments (n=3 animals), we performed online analysis to determine spontaneous PCs and used the leading PCs as design templates for making optogenetic stimuli (Fig. 4H). Across our dataset, we found that stimulus alignment with spontaneous activity was significantly correlated with the trial-to-trial reliability (Fig. 4I; r=0.53, p<0.001, Pearson’s correlation) and similarity to stimulus (Fig 4J; r=0.62, p<0.001), consistent with model predictions. Alignment with dominant spontaneous was also able to explain the degree of response stability for long-duration stimuli, with intra-trial stability and the similarity with the stimulus being both positively correlated with stimulus overlap with spontaneous activity (Fig. 4K,L. Intra-trial stability: r=0.43, p=0.03; Similarity to stimuli: r=0.62, p<0.001, Pearson’s correlation). Taken together, these results show that the degree of input-recurrent alignment determines the degree of selective amplification performed by millimeter-scale cortical networks.

## Discussion

Understanding how millimeter-scale cortical networks process information requires identifying both the nature of selective amplification they support and the input patterns that most effectively engage them. Using targeted optogenetic stimulation in columnar ferret V1, we directly tested model predictions and uncovered core principles governing recurrent network amplification at millimeter-scale. Amplification in the cortex is highly selective and expressed most strongly in the reliability and input-similarity of evoked responses rather than in their amplitude. The degree of amplification is pattern-dependent and increases with the similarity between the stimulation and the distributed modular patterns of the leading principal components of spontaneous activity. Our model offers a mechanistic explanation of these observations by showing that these components are tightly linked to the dominant modes of recurrent connectivity, indicating that selective amplification critically depends on the alignment between external input and recurrent network structure. Input-recurrent alignment thus provides a unifying framework for explaining stimulus-selective amplification within millimeter-scale cortical networks.

Our results identify response reliability – both across trials and during sustained stimulation - as the hallmark of selective amplification in millimetre-scale cortical networks. Classical linear network models predict that the primary effect of aligned input is to produce large response amplitudes (*27, 28*) (Fig. S2), with increased reliability emerging as a consequence of this amplitude effect (*29*). By contrast, our nonlinear model shows that aligned inputs increase reliability with minimal change in response amplitude (Fig. 1F, Fig. S3). This near invariance appears to be a consequence of the rectifying nonlinearity, which stabilizes activity by enforcing a lower bound of zero on firing rates. Consistent with these predictions, our *in vivo* data does not reveal any systematic amplitude effects. Together, these findings argue that recurrent interactions primarily stabilize evoked activity when input aligns with the network’s recurrent structure, rather than enhancing its magnitude.

Our empirical results reveal a striking degree of selectivity, with some input patterns engaging the network far more effectively than others, even when matched for spatial wavelength. Our model attributes this selectivity to local excitation/lateral inhibition (LE/LI) recurrent connectivity that is spatially heterogenous. A circuit motif implementing LE/LI interactions has been found to dominate activity patterns in developing ferret V1 (*25, 26*). LE/LI interactions naturally generate modular activity patterns at the columnar scale and define the characteristic spatial wavelength of amplifiable patterns(*31–33*), features supported by the wavelength-dependent amplification of random inputs seen in our data (Fig. S7). However, owing to the high degree of degeneracy in such spatially homogenous networks, LE/LI alone cannot account for the strong selectivity we observe in our experiments (Fig. S1). Instead, even a moderate degree of symmetry breaking heterogeneity renders amplification in LE/LI networks highly selective (Fig. 1D-F, Fig. 4B), presumably by creating privileged pathways through which specific patterns engage the network more effectively (*26, 33*). Because heterogeneity is a natural consequence of cortical wiring variability, such selective network amplification is therefore expected to be a general feature of columnar cortical networks. Our modeling and *in vivo* data together thus indicate a significant contribution from columnar-scale heterogeneity with local variations in connectivity biasing amplification toward specific pattern layouts, showing that together LE/LI interactions and intrinsic heterogeneity shape the selective amplification landscape.

Our findings suggest a general principle of cortical processing at the millimeter scale–the scale at which perception and behavior engage distributed population activity (*34–39*): that inputs aligning to prevalent spontaneous activity modes yield reliable and stable representations. V1 exemplifies this principle; the two leading spontaneous principal components are closely matched to the orientation preference map in the experienced cortex (*26, 29, 40*). This principle may in fact be shared across cortical areas as similar modular patterns of spontaneous activity occur in both sensory and association cortices (*22, 23*), suggesting that alignment between inputs and intrinsic network modes may represent a widespread cortical strategy.

This insight has important practical implications for neural prosthetics and brain-computer interfaces, where control of local population dynamics remains a major challenge, as it suggests an applicable strategy for identifying amplifiable input patterns directly from spontaneous activity. In fact, stimulation strategies guided by spontaneous activity structure improve control efficacy, predicting perturbation outcomes and enhancing pattern discrimination (*41, 42*), consistent with the selectivity of network amplification shown here. Likewise, manipulations targeting within-manifold modes yield more effective control than out-of-manifold perturbations (*43–46*). Together with our results, this suggests that incorporating the selective amplification of specific activity patterns may help guide next-generation neural interfaces.

Finally, our results also offer a coherent framework for understanding developmental changes in cortical function. At eye opening, visually evoked activity is initially unreliable, before becoming more reliable and stable with visual experience (*20, 29, 47*). Our results suggest that early sensory inputs are poorly aligned with pre-existing recurrent circuits and therefore cannot fully recruit recurrent amplification mechanisms (*20, 29*). Through activity-dependent plasticity, inputs that drive more reliable responses become preferentially reinforced, allowing recurrent amplification of aligned inputs to stabilize emerging sensory representations (*6, 29*) Thus, our findings suggest a bidirectional refinement process where recurrent network interactions shapes inputs, and those inputs simultaneously sculpt the network’s connectivity, offering a solution to the paradox of how reliable sensory representations emerge from initially noisy and immature circuits. Such a framework is consistent with the observed convergence of spontaneous and visually evoked activity onto a shared low-dimensional space during development (*29*). Importantly, this alignment process could operate largely independently of feedforward tuning changes (*20*), providing a complementary mechanism for the emergence of reliable representations. This framework thus offers a lens for evaluating models of cortical development and organization (*21, 48–50*) and motivates longitudinal optogenetic perturbation experiments to causally trace how specific input patterns drive circuit refinement over time.

## Acknowledgments

The authors wish to thank Nic Glewwe, Matt Paruzynski, Jack Kapler, and Sophie Bowman for surgical assistance, Deano Farinella and Harishankar Jayakumar for their optical expertise in designing and maintaining our microscope, and members of the Smith and Kaschube labs for helpful discussions. All viral vectors used in this study were generated by the University of Minnesota Viral Vector and Cloning Core (Minneapolis, MN). This work was supported by the resources and staff at the University of Minnesota University Imaging Centers (SCR 020997).

## Funding

National Eye Institute (NIH R01EY030893; GBS)

National Institute for Neurological Disorders and Stroke (R01NS147764; GBS)

National Institute of Mental Health (MH115688 T32; HNM)

National Science Foundation (IIS-2011542; GBS)

Whitehall Foundation (2018-05-57; GBS)

Bundesministerium für Bildung und Forschung (BMBF 01GQ2002; MK)

LOEWE Research Cluster CMMS (ST, MK)

## Author contributions

Conceptualization: HNM, ST, MK, GBS

Methodology: HNM, ST

Investigation: HNM, ST

Analysis: HNM, ST

Visualization: HNM, ST

Funding acquisition: GBS, MK

Project administration: GBS, MK

Supervision: GBS, MK

Writing – original draft: HNM, ST, MK, GBS

Writing – review & editing: HNM, ST, MK, GBS

## Competing interests

The authors declare that they have no competing interests.

## Data and code availability

All source data for figures are provided with this paper. Due to size limitations, all original raw data (imaging files) are available from the corresponding author upon request. Custom code for computational model simulations will be made available on github at the time of this paper’s publication.

## Supplementary Materials

Materials and Methods

Figs. S1 to S7

References (*1–54*)

## Materials and Methods

### Animals

All experimental procedures were approved by the University of Minnesota Institutional Animal Care and Use Committee and were performed in accordance with guidelines from the US National Institutes of Health. We obtained 9 male and female ferret kits from Marshall Farms and housed them with jills on a 16 h light/8 h dark cycle. All imaging was done in young animals (postnatal day (P)23–29) prior to eye-opening (typically P31 to P35 in ferrets). No statistical methods were used to predetermine sample sizes, but our sample sizes are similar to those reported in previous publications (*17, 26, 47*).

### Viral injection

Viral injections were performed as previously described (*51*) and were consistent with prior work (*25, 52*). Briefly, we microinjected a 1:1 ratio of AAV1.hSyn.GCaMP6s.WPRE.SV40 (Addgene) and the somatically targeted AAV1.hSyn.ChrimsonR.mRuby2.ST (University of Minnesota Viral Vector and Cloning Core) into layer 2/3 of the primary visual cortex at P10–15 approximately 10–15 days before imaging experiments. Anesthesia was induced with isoflurane (3.5–4%) and maintained with isoflurane (1–1.5%). Buprenorphine (0.01 mg/kg) and glycopyrrolate (0.01 mg/kg) were administered, as well as 1:1 lidocaine/bupivacaine at the site of incision. Animal temperature was maintained at approximately 37^°^C with a water pump heat therapy pad (Adroit Medical HTP-1500, Parkland Scientific). Animals were also mechanically ventilated and both heart rate and end-tidal CO2 were monitored throughout the surgery. Using aseptic surgical technique, skin and muscle overlying the skull above the visual cortex were retracted. To maximize the area of ChrimsonR expression, two small burr holes placed 1.5-2 mm apart were made with a handheld drill (Fordom Electric Co.). Approximately 1 *μ*l of virus contained in a pulled-glass pipette was pressure injected into the cortex at two depths (~200 *μ*m and 400 µm below the surface) at each of the craniotomy sites over 20 min using a Nanoject-III (World Precision Instruments). The craniotomies were filled with 2% agarose and sealed with a thin sterile plastic film to prevent dural adhesion, before suturing the muscle and skin.

### Cranial window surgery

On the experiment day, a cranial window was surgically implanted over the imaging area. Atropine (0.2 mg/kg) was injected subcutaneously and then ferrets were anesthetized with 3-4% isoflurane. Animals were placed on a feedback-controlled heating pad to maintain an internal temperature of 37–38^°^C. Animals were intubated and ventilated, and isoflurane was delivered between 1% and 2% throughout the surgical procedure to maintain a surgical plane of anesthesia. An intraparietal catheter was placed to deliver fluids. EKG, end-tidal CO2, and internal temperature were continuously monitored during the procedure and subsequent imaging session. The scalp was retracted and a custom titanium headplate adhered to the skull using C&B Metabond (Parkell). A 6– 7 mm craniotomy was performed at the viral injection site and the dura retracted to reveal the cortex. One 4 mm cover glass (round, #1.5 thickness, Electron Microscopy Sciences) was adhered to the bottom of a custom titanium insert and placed onto the brain to gently compress the underlying cortex and dampen biological motion during imaging. The cranial window was hermetically sealed using a stainless-steel retaining ring (5/16-in. internal retaining ring, McMaster-Carr). Upon completion of the surgical procedure, isoflurane was gradually reduced (0.6–0.9%) and then vecuronium bromide (0.4 mg/kg/hr) mixed in an LRS 5% Dextrose solution was delivered IP to reduce motion and prevent spontaneous respiration.

### Wide-field epifluorescence and optogenetic stimulation

Imaging and optogenetic stimulation were performed using a custom-built opto-macroscope (*24*), providing artifact-free simultaneous imaging of GCaMP signals and optogenetic stimulation. Imaging was performed using a sCMOS camera (Prime BSI express, Teledyne) at 15 Hz with 2×2 binning and additional offline 2×2 binning to yield 512×512 pixels. Optogenetic stimulation is provided by a 590nm continuous laser (Coherent MX590–1000 STM OPSLaser-Diode), and optogenetic stimuli were projected onto the surface of the brain using a digital micromirror device (DMD) pattern illuminator (Mightex POLYGON1000-DL).

#### Imaging spontaneous activity

At the onset of experiments, prior to optogenetic stimulation, approximately 30-60 minutes of baseline spontaneous activity was captured in 10 minute imaging sessions, with the animal sitting in a darkened room facing an LCD monitor displaying a black screen.

### Design of spatially structured optogenetic stimuli

For patterned stimulation, binary optogenetic masks were made using Polyscan software (Mightex) or Matlab (Mathworks). Stimulation patterns were based on endogenous patterns of activity observed in spontaneous activity, or random patterns (see below). To test the specificity of opto-evoked responses, for each experimental condition at least 3 different spatial patterns were created.

#### Endogenous stimulation patterns

To approximate endogenous network patterns, we created optogenetic masks based on spontaneous correlation patterns, which are robust to event number inclusion criteria and stable over several hours (*26*). All optogenetic stimulation took place within this time window. Online detection of spontaneous events was done by using the Matlab function ‘findpeaks’ to find events in the ΔF/F trace (averaged over the full Region of Interest (ROI) encompassing the area of GCaMP expression) that were greater than 1 standard deviation over the mean.

These events were spatially filtered (Gaussian highpass filter, *σ*_high_=195 *μ*m) yielding spontaneous activity patterns 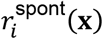 for event *i* and location (**x**). From these we calculated the correlation patterns *C*(**s, x**), representing the pairwise Pearson’s correlation between the activity at all locations **x** within the ROI and a seed point **s**:

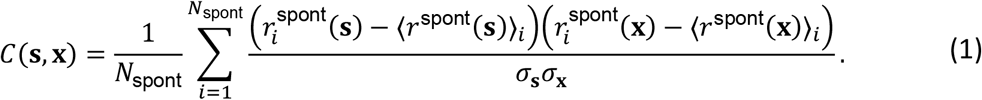

Here, *i* is the pattern index (*i* = 1 … *N*_spont_), ⟨⋅⟩_*i*_ denotes the average over all *N* spontaneous patterns (e.g., 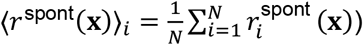, and *σ*_x_ is the standard deviation of activity at location **x** over all *N* patterns.

We arbitrarily selected seed points that showed modular patterns with multiple positively correlated domains that extended over millimeters of cortical surface, aiming to select patterns that were spatially distinct (see Figure S3 for examples). Opto-stimulus masks were generated by either thresholding all pixels with correlations greater than 0.15-0.2, or by manually drawing regions around positively correlated domains.

#### Random stimulation patterns

To produce random stimulation patterns (experimental data in Figures 2G-J and 3E-H), optogenetic masks were made by bandpass filtering white noise (256 x 256 pixels) in the frequency domain using a 2D Fermi filter (temperature parameter T=0.05), as described in (*25*). Filter cutoffs were first fixed in the spatial domain, with cutoff-frequencies defined by the minimal and maximal wavelengths, *d*_1_ and *d*_2_ (*d*_1_ =40, *d*_2_ =42 pixels, corresponding to frequency cutoffs of 6.1 and 6.4 cycles per image width for N=256 pixels), which when multiplied by the spatial resolution of the imaging camera resulted in real-space wavelengths of 606 µm and 637 µm (for a resolution of 15.16 µm per pixel). This resolution varied slightly across animals (between 14.7 and 15.9 µm per pixel). These parameters were chosen to be similar to the network’s intrinsic characteristic wavelength, yielding a measured wavelength of approximately 0.7 mm, which is consistent with the wavelength of spontaneous activity (Figure S7).

For Figure 2K-M and Figure 4I-L, we pooled together experiments with stimuli generated using a broader spectrum filter (*d*_1_ =30, *d*_2_ =50 pixels, 454-768um), but similar measured wavelength (0.63 mm). Additionally, we analyzed responses to random stimuli of varying spatial scales (Figure S7), where the fixed digital bandpass filter cutoffs were systematically varied from 20 to 62 pixels (step size=10 pixels, bandwidth=2 pixels), corresponding to a physical spatial sweep spanning from approximately 303 µm to 939 µm (wavelength approx. 0.48-1.10mm).

To make a binary optogenetic mask from these patterns, for all patterns we set a threshold at the 68th percentile of the pixel values, producing isolated blobs with a specific wavelength.

#### Principal components of spontaneous activity

For a subset of experiments, we also designed stimulation masks based on the principal components of spontaneous activity (n=3 animals). Principal component (PC) analysis was done on the same filtered spontaneous events, using Matlab’s built in ‘pca’ function. Optogenetic masks based on these components were drawn in a similar manner as the endogenous patterns, by manually drawing or thresholding the pixels (> or < 0) with weights of the same sign. For each animal, 5-6 patterns were drawn from a variety of principal components, targeting both leading and higher order PCs (1-20), but primarily focusing on the most dominant PCs (1-5). The opto-stimulus masks drawn from the online analysis of PCs were similar to the principal components of spontaneous activity after full preprocessing (see below), showing that our rapid online analysis accurately captured the spatial arrangement of the linear dimensions that carry the majority of variance in our dataset of spontaneous activity. Stimulus PC index shown Fig. 4 indicates the most correlated spontaneous PC (after full preprocessing) for the stimulus.

### Stimulation protocol

As with spontaneous activity, optogenetic stimulation was similarly delivered in the absence of visual stimuli, and the animal’s eyes were shielded from the stimulation laser to prevent indirect stimulation of the retina. The sequence of opto-stimulus masks and timing of stimulation was controlled using Polyscan software (Mightex), For all experiments, the power density of photostimulation was approximately 10mW/mm^2^.

Optogenetic stimuli were delivered for 1 second duration and with a 5 second inter-stimulus interval (N=40 repetitions/pattern). Different patterns were interleaved within a given experiment. For experiments where we assessed responses to sustained stimuli (Fig. 3), stimulation patterns were delivered for 5 seconds with a 5 second inter-stimulus interval.

## Data analysis

### Signal extraction for wide-field epifluorescence imaging

Image series were motion corrected using rigid alignment and a region of interest (ROI) was manually drawn around the cortical region of GCaMP expression, excluding major blood vessels. The baseline fluorescence (F_0_) for each pixel was obtained by estimating approximately the 15^th^ percentile of activity within a sliding bin over time, which was calculated by applying a rank-order filter to the raw fluorescence trace F with a rank 70 samples and a time window of 30 s (451 samples). The rank and time window were chosen such that the baseline faithfully followed the slow trend of the fluorescence activity. The baseline-corrected activity was calculated as

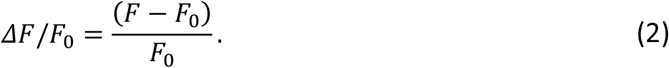

### Event detection and preprocessing

Detection of spontaneously active events was performed as previously described (*26, 52*). Briefly, we first determined active pixels on each frame using a pixelwise threshold set to 5 s.d. above each pixel’s mean value across time. Active pixels not part of a contiguous active region of at least 0.01 mm^2^ were considered ‘inactive’ for the purpose of event detection. Active frames were taken as frames with a spatially extended pattern of activity (>80% of pixels were active). Consecutive active frames were combined into a single event starting with the first high-activity frame and then either ending with the last high-activity frame or, if present, an activity frame defining a local minimum in the fluorescence activity. To assess the spatial pattern of an event, we extracted the maximally active frame for each event, defined as the frame with the highest activity averaged across the ROI.

Opto-evoked evoked events were defined as the single frame response at opto-stimulus offset. To normalize for differences in baseline fluorescence between spontaneous and opto-evoked events, we calculated the mean for all opto-evoked responses across all stimuli (including endogenous and random) within an experiment to determine the ‘global mean’ and then subtracted this global mean from all evoked responses. Then, all events–spontaneous and opto-evoked–were spatially filtered with a bandpass filter using Gaussian filter kernels (*σ*_low_=25 *μ*m, *σ*_high_=195 *μ*m). For long duration experiments (Fig. 3), all frames over the full duration of the trial were spatially filtered with a bandpass filter, however, we did not subtract a global mean. Additionally, global mean was not subtracted for analyses of activity amplitude (Fig 2K, S6).

## Computational modeling

To model selective amplification in a modular cortical recurrent network, we considered a nonlinear firing rate model capable of generating modular activity and probed it with inputs of different spatial structure. The model employs local-excitation/lateral-inhibition (LE/LI) connectivity with moderate, biologically realistic spatial heterogeneity, following approaches that have shown to successfully recapitulate spontaneous dynamics in developing ferret visual cortex (*20, 25, 26*). Our goal was a mechanistic understanding of selective amplification in such nonlinear recurrent network.

### Network model

Building on a standard framework for nonlinear rate models, the dynamics of the firing rate *r*(**x**, *t*) for a unit at position **x** on a two-dimensional grid are described by

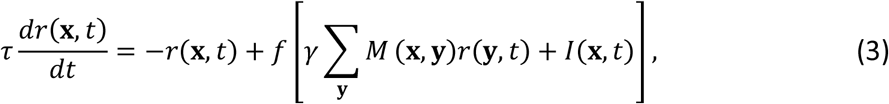

where *τ* is the neuronal time constant (set to 1 a.u.), *f*(*x*) = max(0, *x*) is a rectifying non-linear transfer function (ReLU) ensuring non-negative firing rates, *γ* is the global strength of recurrent interactions, *M*(**x, y**) defines the effective recurrent connectivity, and *I*(**x**, *t*) denotes the external input (specified below). This formulation follows a classic one-population Wilson-Cowan-type approach (*53, 54*), which combines excitatory and inhibitory contributions into the effective connectivity **M**. This network was implemented on a grid with 60 × 60 rate units.

Following (*25*), the recurrent connectivity takes the form

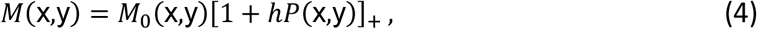

where *M*_0_(**x, y**) is a spatially homogeneous, isotropic connectivity that exhibits a LE/LI organization to ensure the network produces modular activity, and *P*(*x, y*) is a perturbation to introduce moderate heterogeneity to the network. The parameter *h* controls the strength of the heterogeneity, and [*x*]_+_ = max(0, *x*) denotes rectification.

The connectivity matrix *M*_0_ (**x, y**) was implemented by a difference-of-Gaussians,

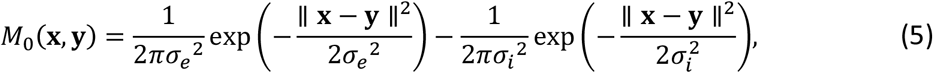

where *σ*_*i*_ and *σ*_*e*_ control the spatial range of lateral inhibition and local excitation, respectively. By choosing their ratio to be larger than one, i.e. *κ* = *σ*_*i*_/*σ*_*e*_ > 1, this connectivity has a LE/LI-organization. Moreover, since it is homogenous in space, the eigenvectors of *M*_0_ are plane waves, and the spectrum is peaked at a wavenumber *k*_0_ = 2*π*/*Λ* with the typical spatial scale *Λ* given by

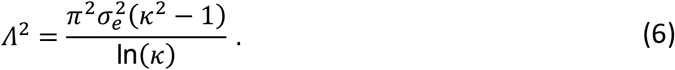

We used *σ*_*e*_ = 1.8 and *σ*_*i*_ = 3.6, implying a wavelength of approximately *Λ* ~ 12 (in Δ*x*, the spacing between units).

To add a biologically realistic, yet generic spatial heterogeneity to this connectivity, we constructed the perturbation field *P*(**x, y**) as a spatially correlated Gaussian random field as follows: First, we initialized a spatial field of independent and identically distributed Gaussian white noise *η* ~ 𝒩(0,1). This noise field was then filtered using a difference-of-Gaussians filter defined by two spatial scales, *σ*_*P*1_ and *σ*_*P*2_, set to 2 and 6, respectively. The resulting field was normalized to have zero mean and unit variance to ensure consistent perturbation strength. Morevoer, we rescaled the recurrent connections to each network unit to have zero mean and uniform variance, consistent with homeostatic mechanisms balancing the input between excitation and inhibition equally across cortex. To achieve this, we first rescaled for each location **x** all negative recurrent weights from **y** to **x**, such that the sum of recurrent weights to location **x** equals zero. Next, we normalized the connections so that the standard deviation *σ*_**y**_ across all recurrent connections to location **x** is 1. Finally, to simplify further analysis and allow interpretation of the parameter *γ*, the connectivity matrix **M** - containing the connection strengths between all units of our network - was scaled such that the real part of its maximal eigenvalue is equal to 1. Such spatial heterogeneity captures possibly variations of connectivity across the cortex and was found in previous work to be a critical parameter in recapitulating the structure of *in vivo* activity (*20, 25, 26*).

### Linear network and SVD analysis

To formalize how our recurrent network preferentially amplifies certain input patterns (Fig. 1), we considered the linearized version of the network. Let the vector **r**(*t*) ∈ ℝ^*N*^ describe the activity of all *N* units in the network (representing the flattened vector of rates *r*(**x**, *t*) of all units in the two-dimensional grid defined above), then its rate of change is described by

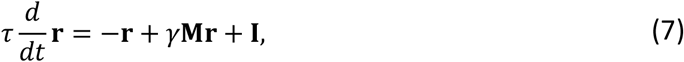

where *γ* < 1 to ensure its dynamics are in a subcritical regime and stable, **M** is now a *N* × *N* recurrent connectivity matrix, and **I** the input vector of length *N*. The steady-state response **r**_*ss*_ to a constant input **I** is found by setting 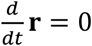, which yields

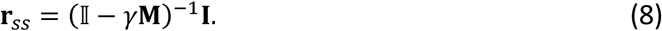

This equation defines a linear mapping from input **I** to response **r**_*ss*_ = **A I** via an effective amplification matrix (also called resolvent matrix)

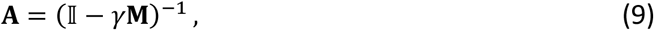

which encapsulates how the recurrent connectivity **M** transforms, and possibly amplifies, feedforward inputs in this linear network.

Applying Singular Value Decomposition (SVD), we can write the resolvent matrix as a sum of rank one connectivity components,

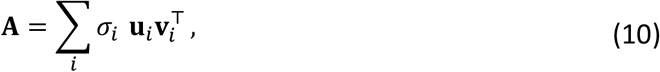

where *σ*_*i*_ ≥ 0 are the singular values, representing the amplification gain, **v**_*i*_ are the right singular vectors, and **u**_*i*_ are the left singular vectors. The sets of right singular vectors **v**_*i*_ and left singular vectors **u**_*i*_ are each orthonormal, forming independent bases for the input and output spaces, respectively. They are vectors of length *N* and correspond to two-dimensional spatial patterns when reshaped to the geometry of the two-dimensional network grid. We term **v**_*i*_ the IN connectivity modes (as they reveal the spatial configurations of inputs the network is sensitive to) and **u**_*i*_ the OUT connectivity modes (as they reveal the spatial patterns of responses elicited by these inputs). An input perfectly aligned with the leading IN connectivity mode, **I** = *a* ⋅ **v**_1_ (where *a* is a scalar amplitude), yields a maximally amplified response **r**_*ss*_ = *a* ⋅ *σ*_1_ ⋅ **u**_1_ aligned with the leading OUT connectivity mode. An input aligned with the *k*-th IN mode (**I** ∝ **v**_*k*_) elicits a response proportional to the *k*-th OUT mode (**r**_*ss*_ ∝ **u**_*k*_), with amplification *σ*_*k*_. Due to the asymmetry of **M**, the IN and OUT modes in general exhibit a distinct spatial structure.

### Network input

For the primary analyses in this study (Fig. 1,2,4), we investigated the network’s steady state response to static spatial inputs. The network’s recurrent dynamics (Eq. 3) thus evolve in response to a static input pattern *I*(**x**). The general form for this input is

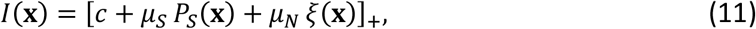

where *c* is the constant baseline input, *P*_*S*_(**x**) is a spatially structured component, with zero mean, scaled by *μ*_*S*_, and *ξ*(**x**) is a white noise process (𝒩(0,1)), scaled by *μ*_*N*_. We set *c* = 1 to represent tonic background input, consistent with the ongoing synaptic drive cortical neurons receive in vivo. This ensures that all units initially receive suprathreshold input and contribute to the recurrent dynamics.

The parameters *μ*_*S*_ and *μ*_*N*_ control the relative strengths of the structured stimulus and unstructured noise, respectively, and their ratio determines an overall level of signal-to-noise-ratio in the input. We simulated two primary input conditions based on this general form: unstructured noise input (*μ*_*S*_ = 0) to simulate spontaneous activity (following (*20, 25, 26*)) and structured optogenetic stimulation (*μ*_*S*_ > 0).

#### Spontaneous activity

To simulate the network’s spontaneous activity, we set the structured stimulus strength *μ*_*S*_ = 0 in Eq. (11). The input for a single spontaneous pattern *i* is thus

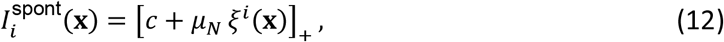

for which we simulated a set of *N* = 200 spontaneous activity patterns 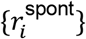 with different noise realizations *i* to approximate the endogenous activity manifold. This sample size is sufficient to stably estimate the dominant correlations in the spontaneous correlation structure (see (*26*)).

#### Structured input patterns

To test how the network responds to different spatial structures in the input and to examine which inputs are selectively amplified, we used the structured input condition, defined by *μ*_*S*_ > 0. The spatially structured components *P*_*S*_(**x**) were drawn from several distinct sources, which we describe in the following.

#### Connectivity modes

To test the responses of the full nonlinear recurrent network to inputs matching its IN connectivity modes – representing the optimal spatial configurations to drive recurrent amplification in the linearized network (see SVD analysis above and Fig. 1) – we computed the resolvent matrix **A** in a regime of strong net connectivity (*γ* = 0.99). The SVD of **A** yields **A** = **USV**^⊤^, where the columns of **U** and **V** contain the OUT and IN connectivity modes, respectively, and the diagonal entries of **S** contain the singular values. To obtain the spatial stimulation patterns *P*_*i*_(**x**), we selected the IN connectivity modes **v**_*i*_ (columns of **V**) and reshaped them into the two-dimensional spatial grid of the nonlinear network. For the primary results (e.g., Fig. 1F), we simulated responses for every second mode starting from the leading one (index 1, 3, …). For the robustness controls involving extensive parameter sweeps (Fig. SI2), we utilized a coarser sampling of every third mode (index 1, 4, 7, …) to maintain computational efficiency.

#### Spontaneous correlation patterns

To test responses to ‘endogenous’ stimulation patterns in the nonlinear model (Fig. 2), we used correlation patterns of spontaneous activity as structured inputs, analogous to the endogenous stimulation patterns in our *in vivo* experiments above. These correlation patterns summarize typically co-active domains with the seed point and were computed using the network’s simulated patterns of spontaneous activity (*N*_*spont*_ = 200 spontaneous activity patterns). As for the experimental data, we calculated correlations patterns *C*(**s, x**) (see Eq. 1), and then randomly selected a set of *k* seed locations {**s**_*j*_} (for *j* = 1… *k*) to define the corresponding input patterns *P*_*S*_ = *C*(**s**_*j*_, **x**).

#### Random bandpass filtered patterns

To investigate whether network amplification systematically varies with the characteristic spatial wavelength of input patterns relative to the network’s intrinsic characteristic wavelength (see Fig. SI 7), we generated random patterns with controlled range of spatial frequencies across various scales by bandpass-filtering spatial white noise using a hard cutoff filter in the frequency domain (following the procedure in (*25*)). First, to compare endogenous pattens to random patterns we matched their wavelength to the network’s characteristic wavelength (*d*_1_ = 10, *d*_2_ = 14, matching the intrinsic wavelength of *Λ* = 12, all in units of unit spacing Δ*x*). Second, to systematically investigate the dependency on spatial scale, we generated bandpass patterns by varying the center wavelength *d*_*c*_ from 6 to 20Δ*x* in 15 linearly spaced steps. For each center wavelength, we maintained a constant bandwidth of 3Δ*x*, setting the cutoffs as *d*_1_ = *d*_*c*_ − 1.5 and *d*_2_ = *d*_*c*_ + 1.5 Δ*x*. For each wavelength range, independent patterns were generated from different noise realizations.

#### Spontaneous principal components

To test whether stimulus alignment with the principal components (PCs) of spontaneous activity predict network amplification (see Fig. 4), we computed Principal Component Analysis (PCA) on simulated spontaneous network activity (see above). To test the dependency on the order of the spontaneous PCs **p**_*i*_, we chose the stimulation patterns directly as principal component *P*_*S*_(**x**) = **p**_*i*_. For the primary results (Fig. 4), we tested every second PC starting from the principal one (index 1, 3, …). For the robustness controls involving extensive parameter sweeps (Fig. S3), we utilized a coarser sampling of every third PC (index 1, 4, 7, …) to maintain computational efficiency.

#### Patterns sampled from spontaneous manifold

Additionally, to explore a more diverse relationship between network amplification and alignment with spontaneous activity (Fig. 4 D-G), we constructed stimulus patterns as a weighted linear combination

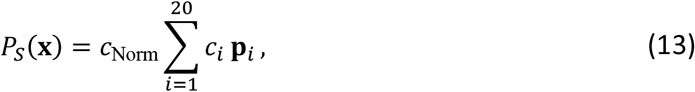

of the leading 20 most dominant PCs **p**_*i*_, where *c*_Norm_ is a normalization factor applied to scale all generated patterns to have the same root-mean-square (RMS) contrast. The coefficients *c*_*i*_ were drawn from a zero-mean Gaussian distribution, 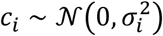, with the variance decaying as 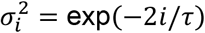 with component rank *i* to have the generated stimuli overlapping more strongly with the leading PCs. Here, *τ* is a decay constant that sets the steepness of the falloff and determines the dimensionality of the space we draw patterns from (see (*29*)). A falloff *τ* = 20 was chosen so that it encompasses the space of spontaneous activity and ensures we are also probing the network with unaligned stimuli.

#### Binarization and trial structure

To mimic 1-photon optogenetic stimulation, a stimulation pattern was binarized and thresholded as

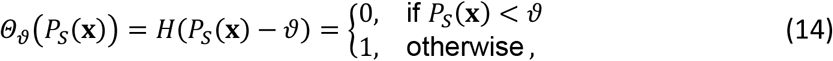

where the threshold ϑ was set at the 68th percentile of the template pattern’s values, *P*_68_[*P*_*S*_(**x**)]. This threshold was set to replicate the stimulus structure used in optogenetic stimulation (*24, 25*); it is approximately one standard deviation above the mean for normally distributed input strength, thereby preserving the topological features of the source domains. Here, following Eq. (11), the same structured pattern Θ_ϑ_(*P*_*S*_(**x**)) is presented on every trial, while a different noise field *ξ*^*i*^(**x**) is drawn for each trial *i* to model variability, resulting in response patterns 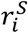 used for subsequent analysis.

### Sustained simulation

To model network dynamics in response to sustained inputs (Fig. 3B,C), we used a static base pattern with an additional time-dependent noise component with a controlled temporal correlation. The total input was thus defined by

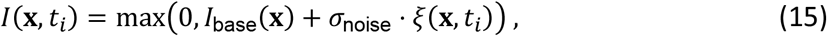

where the base pattern *I*_base_(**x**) follows Eq. (11), combining a baseline, structured stimulus component and static trial noise; *σ*_noise_ controls the relative strength of the fluctuating component, and *ξ*(**x**, *t*_*i*_) is a dynamic, spatially-uncorrelated, but temporally-correlated noise field. This correlated noise *ξ*(**x**, *t*_*i*_) was generated as a first-order autoregressive (AR(1)) process. At each time step *t*_*i*_, a field of independent white noise *η*(**x**, *t*_*i*_) ~ 𝒩(0,1) was drawn. The correlated noise *ξ* was then updated according to

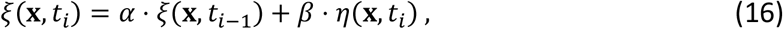

where *t*he coefficients *α* and *β* are determined by the desired noise timescale, *τ*_noise_, and the simulation time step, Δ*t*. The “memory” coefficient *α* = exp(−Δ*t*/*τ*_noise_) defines the decay of the correlation, and the “innovation” coefficient 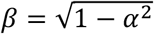 scales the new white noise to ensure the resulting process *ξ* has a stationary variance of 1. We used a noise timescale of *τ*_noise_ = 20 (in simulation time units) to emulate the slow fluctuations of cortical response patterns rather than fast synaptic noise.

### Implementation

We implemented the network with 60 × 60 units placed on a regularly-spaced grid and periodic boundary conditions with custom-written code in Python, using the tensorflow package for calculations on an *M*4 apple chip. We numerically integrated the dynamics (Eq. 3) using a fourth-order Runge-Kutta method from *t* = 0 to *t* = *T* using a time step *dt* = 0.15*τ*, with initial conditions *r*(**x**, *t* = 0) = 0. The integration time was set to *T* = 500 (unless noted otherwise) to ensure that activity patterns reached a near steady state under all used input conditions.

Similar to the empirical data, for each instance of the connectivity, we simulated responses for *N*_trial_ = 10 trials for each stimulus pattern (unless noted otherwise). These different trials used different instances of the noise process *ξ*(**x**) (see Eq. 11). The AR(1) noise process for sustained stimulation (Fig. 3) was implemented using a digital filter (*scipy.signal.lfilter*).

### Model parameters and validation

The model’s free parameters (including recurrent strength *γ*, heterogeneity strength *h*, input offset *c*, and noise amplitude *μ*_*N*_) were selected to ensure the model operated in a realistic dynamical regime, consistent with experimental observations from developing ferret visual cortex. We systematically varied these parameters for the simulation of spontaneous activity. The final parameter set was chosen by validating that the resulting spontaneous activity met three key criteria: the dimensionality of the activity fell in the range of literature values; the activity exhibited spatial modularity levels comparable to *in vivo* data; and the activity patterns were visually similar to characteristic epifluorescence recordings.

This validated set of parameters was then fixed and used for all simulations described in this study (see Table 1 for an overview). The stimulus amplitude *μ*_*S*_ was chosen to be small relative to 1, allowing the recurrent dynamics to strongly influence the responses. Parameters that we chose for the specific figures are shown in Table 2.

Crucially, our primary findings are not dependent on this specific parameter combination. As shown in Fig. SI 2, the qualitative results regarding the effects of alignment remain robust against variations in recurrent strength *γ* and input signal-to-noise ratio over a significant range. The chosen parameter set is a representative point within this regime.

#### Reduced model configurations

To control for the role of two central model features, heterogeneity and nonlinearity, we studied two model variants with reduced features in Fig. S1 and S2, respectively.

#### Homogeneous network (Figure S1)

To determine whether the selectivity of amplification, the network’s preference for specific patterns over others of the same wavelength, arises from recurrent heterogeneity, we simulated a homogeneous network (*h* = 0). In this condition, the connectivity matrix **M** contains only the distance-dependent LE/LI component. To probe the network’s selectivity, the additive noise *ξ*(**x**) was generated using a bandpass Difference-of-Gaussians filter with spatial scales *σ*_*low*_ = 1 and *σ*_*high*_ = 2 Δ*x*. This produced a noise spectrum that was broader than the network’s intrinsic scale (covering a wider range of frequencies than the specific intrinsic wavelength), testing the network’s ability to selectively amplify specific components purely based on uniform connectivity rules. To minimize defects due to a simulation within a finite-size network, we chose a smaller intrinsic wavelength (*Λ* ~ 10) by setting *σ*_*e*_ = 1.53 and *κ* = 2, and integrated for *T* = 1200*τ*. To ensure the homogeneous network produced smooth activity patterns, the additive noise ξ(x) was bandpass filtered using Gaussian filter kernels (*σ*_*Low*_ = 1Δx and *σ*_*High*_ = 2 Δx). We verified that the selective amplification results in the heterogeneous network are robust to using this bandpass noise instead of white noise.

#### Linear Network (Figure S2)

*To compare our results for the nonlinear network to amplification in a linear network (Fig. SI 1), we directly calculated its steady-state responses using the analytical solution (Eq. 8*). For the linear network, the baseline input was set to c = 0. Because the linear model decomposes the response additively, a nonzero baseline *would* generate a response component that is identical across all stimulus conditions, *which would be* uninformative for comparing stimulus-specific amplification.

#### Parameter overview

Parameters used for the full nonlinear heterogeneous network (studied in Figs. 1-4) are presented in Table 1 unless noted otherwise in Table 2. This configuration combines fixed simulation constants and standard model parameters.

#### Standard Model Configuration

| Parameter | Symbol | Value |
| --- | --- | --- |
| *Spatial & Temporal Resolution* |  |  |
| Grid resolution | $\Delta x$ | 1 (arbitrary units) |
| Integration time step | $dt$ | 0.15 |
| Neuronal time constant | $\tau$ | 1.0 |
| Integration duration (Standard) | $T$ | 500 |
| Integration duration (Sustained Stim.) | $T$ | 900 |
| *Network Structure* |  |  |
| System Size | $N$ | $60 \times 60$ |
| Recurrent strength | $\gamma$ | 1.05 |
| Excitatory spread | $\sigma_e$ | 1.8 |
| Ratio Inhibition/Excitation spread | $\kappa$ | 2 |
| Heterogeneity strength | $h$ | 0.4 |
| Heterogeneity scales | $\sigma_{P1}, \sigma_{P2}$ | 2, 6 |
| Max eigenvalue (normalized) | $\lambda_{max}$ | 1.0 |
| *Input constants* |  |  |
| Baseline tonic input | $c$ | 1.0 |
| Static noise amplitude | $\mu_N$ | 0.2 |
| Stimulus amplitude | $\mu_S$ | 0.06 |
| Stimulus threshold | $\vartheta$ | 68<sup>th</sup> percentile |
| Dynamic noise amplitude | $\sigma_{noise}$ | 0.2 |
| *Input Pattern Generation* |  |  |
| Number of spontaneous patterns | $N_{spont}$ | 200 |
| Intrinsic Wavelength | $\Lambda$ | $\approx 12$ pixels |
| Random Bandpass (control) | $d_1, d_2$ | 10,14 pixels |
| Spontaneous sampling decay | $\tau_{PC}$ | 20 |
| Input correlation timescale | $\tau_{noise}$ | 20 |

| Figure / Analysis | $N$ | $\gamma$ | $\mu_S/\mu_N$ | T | h | Other Deviations / Notes |
| --- | --- | --- | --- | --- | --- | --- |
| Standard Model (Ref) | 60 | 1.05 | 0.3 | 500 | 0.4 | $\Lambda \sim 12$ px |
| Fig. S1 (Homogeneous) | — | — | — | 1200 | 0 | $\sigma_e = 1.53$; Gaussian noise |
| Fig. S2 (Linear) | — | 0.99 | — | — | — | c=0 |
| Fig. S3 (Gamma) | 50 | 0.9 -1.15 | — | — | — |  |
| Fig. S3 (SNR) | 50 | — | 0.1 – 0.9 | — | — |  |
| Fig. S7 (Wavelengths) | — | — | — | — | — | $d_c = 6 - 20$ px (step size 1) ;<br>Bandwidth d = 3 px ($d_c \pm 1.5$) |

Values matching the Standard Model are marked with “—“ in Table 2.

## Analysis

All measures presented in the following are equally applied to the recorded and preprocessed experimental data and modelling results, unless noted otherwise.

### Measures of pattern amplitude

#### Average amplitude within top 10% active pixels

To quantify the amplitude of stimulus evoked responses to a given stimulus pattern *P*_stim_(**x**), we computed the activity within the 10% most active pixels of the resulting activity pattern *r*(**x**).

All measures are computed within a binary Region of Interest (ROI) mask, Ω(**x**). For the computational model, the activity *r*(**x**) represents the steady-state firing rate and the ROI covered the full simulation grid. For the experimental analysis, *r*(**x**) corresponds to the pixel-wise Δ*F*/*F* signal (without band-pass filtering), and the ROI was manually defined to encompass the viral expression zone.

The mean activation *μ*_10%_ within the 10% most active pixels for each event inside the ROI was calculated as

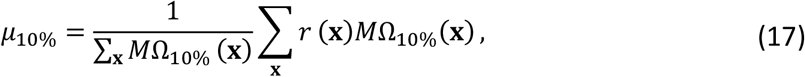

where Ω*M*_10%_(**x**) is a binary mask selecting the top 10% most active locations within the ROI. Specifically, Ω*M*_10%_(**x**) = 1 if the location is within the ROI (Ω(**x**) = 1) and its activity *r*(**x**) exceeds the 90th percentile of all ROI activity values; otherwise Ω*M*_10%_(**x**) = 0. The threshold of 10% was chosen to capture the peak response while ensuring a sufficient number of pixels for stable averaging. We found that this threshold typically captured the active modular domains in each event. All resulting measures were averaged over trials and stimuli, for the respective analyses across stimuli or animals.

#### Alternative measures of response amplitude

To ensure we could capture any modulation of response amplitude, we also quantified responses using two additional metrics: activity within stimulated pixels, and spatial contrast. This multi-metric approach ensures that any modulation of activity, whether affecting only the most active units, localized to the stimulated region, or distributed, would be captured, and allows us to verify that our results are robust to the specific choice of response metric (see comparison in Fig. S2).

#### Stimulated region average amplitude

To quantify the response specifically driven by the input, we calculated the mean activation *μ*_stim_ only within the region directly stimulated, defined as

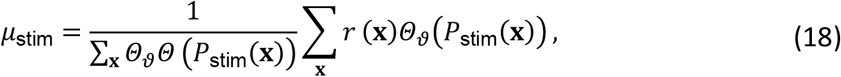

where Θ_ϑ_(*P*_stim_(**x**)) is the binary thresholded stimulus pattern (1 for stimulated locations, 0 otherwise) as described above. The denominator ∑_**x**_ *Θ*_*ϑ*_*Θ* (*P*_stim_(**x**)) is the total number of stimulated locations.

#### Spatial contrast

Finally, to measure the spatial non-uniformity or “contrast” of the activity pattern within the ROI, we computed the spatial standard deviation *σ*_spatial_ for each trial, given by

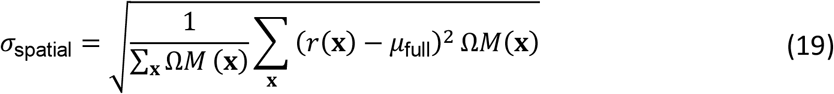

where *r*(**x**) is the activity, Ω(**x**) is the binary ROI mask, and *μ*_full_ is the average activation across the ROI for that trial. This per-trial contrast was then averaged across trials and stimuli.

### Similarity to stimulus

To quantify the similarity between a single-trial evoked response and its corresponding stimulus (as illustrated in Fig. 1F, 2D for the model and Fig. 2L for data), we computed the Pearson correlation between the response pattern and the binarized stimulus pattern. The correlation 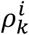 for trial *k* evoked by the *i*-th stimulus pattern is

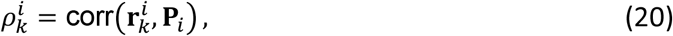

where 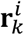 is the response pattern from trial *k* (firing rates for the model; filtered Δ*F*/*F* for the experiment) and **P**_*i*_ is the binarized stimulus pattern (input pattern for the model; projected optogenetic pattern for the experiment), both flattened to a vector. Trial-level correlations were averaged across trials to obtain a stimulus-specific similarity *ρ*^*i*^, and then across stimuli to obtain an animal-level score *ρ*_animal_.

### Trial-to-trial correlation

To measure response reliability (e.g. Fig. 1F, Fig. 2M), we computed the Pearson correlation between all pairs of single-trial response patterns evoked by the same stimulus. For stimulus *i*, the correlation between trials *k* and *l* is

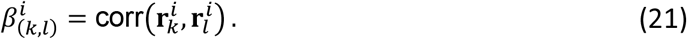

The trial-to-trial reliability *C*_*i*_ was defined as the mean across all *N*_trials_(*N*_trials_ − 1)/2 unique trial pairs.

Controls in the experimental data were estimated by randomly trial shuffling the stimulus IDs and taking the average correlation of event number matched trial pairs, then finding the mean across 100 random shuffle simulations (Fig. SI 5). Additionally, ‘Endo’ evoked trial-to-trial reliability was compared to the naturally occurring similarity across detected spontaneous events (Fig. S5), which was calculated by randomly selecting an event-number-matched subsample of spontaneous and calculating the event-wise Person’s correlation.

### Alignment with connectivity modes

To quantify in the model the degree of alignment of a stimulus input pattern **h** with the network’s recurrent connectivity (e.g. Fig. 2B), we defined a measure of stimulus-network alignment by generalizing an alignment measure previously used for symmetric connectivity (**M** = **M**^*T*^) (*29*). There, the alignment *v* of an input **h** was defined by the quadratic form

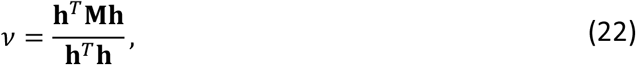

which compares the input’s projection onto the connectivity with its own magnitude. This formulation is equivalent to a sum over the input’s overlap with the network’s eigenvectors weighted by the eigenvalues. Using the spectral decomposition 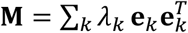 (where *λ*_*k*_ are the (real) eigenvalues and **e**_*k*_ are orthonormal eigenvectors), the numerator becomes **h**^*T*^**Mh** = ∑_*k*_ *λ*_*k*_ (**h** ⋅ **e**_*k*_)^2^. This yields the equivalent form

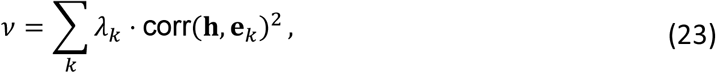

where corr(**h, e**_*k*_)^2^ = (**h** ⋅ **e**_*k*_)^2^/(∥ **h** ∥^2^∥ **e**_*k*_ ∥^2^) = (**h** ⋅ **e**_*k*_)^2^/∥ **h** ∥^2^.

However, this eigen decomposition is not suitable for the asymmetric matrix **M**. We therefore generalized this measure by analyzing the network’s steady-state input-output mapping, described by the resolvent matrix ***A*** = (**I** − **M**)^−1^. Using SVD, as above, provides a set of orthogonal “amplification components,” each defined by an IN-connectivity modes **v**_*k*_ (right singular vector) and an OUT-connectivity modes **u**_*k*_ (left singular vector).

As illustrated in Figure 2B, the overlap between a stimulus **P**_*i*_ and the network’s preferred IN-connectivity modes was quantified by the absolute Pearson’s correlation, |corr(**P**_*i*_, **v**_*k*_)|. To create a scalar alignment metric analogous to Eq. (22), we replaced eigenvalues with singular values and eigenvectors with the input singular vectors, which yielded

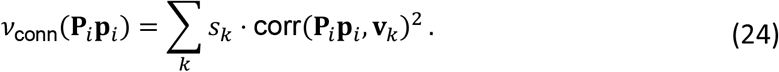

To normalize this measure for binarized stimulus patterns, we first defined a normalization constant *c*_SVD_ based on the first input singular vector **v**_1_, which represents the network’s dominant amplification mode

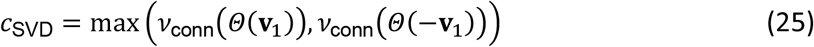

where *v*_conn_(⋅) is the non-normalized alignment from Eq. (24), and Θ is the binarization operator. This constant represents the maximum alignment score achievable by a binarized pattern derived from the primary input mode. This normalization facilitates the comparison of alignment scores across different network realizations and animals and is used for the results presented in Fig. 4. The final normalized alignment of a stimulus pattern **P**_*i*_ to the recurrent connectivity is then given by

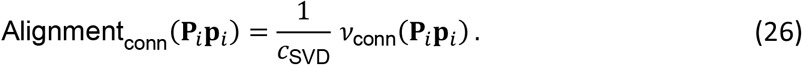

This measure provides a single score for how strongly a binarized stimulus pattern aligns with the network’s high-gain connectivity modes, relative to the maximum possible alignment for a binarized pattern.

### Intra-trial reliability

To test whether the temporal evolution of evoked activity patterns was reliable within a single trial, we computed two distinct measures: one assessing temporal stability (how much the pattern changed from one moment to the next) and one assessing temporal fidelity (how well the pattern matched the stimulus over time).

#### Intra-trial stability (temporal stability)

To measure how much the spatial pattern changed over time (Fig. 3B,C) during a sustained response, we measured the lagged correlation of the spatial pattern at a fixed time lag *τ*_lag_. For a given stimulus *i* and trial *k*, we computed

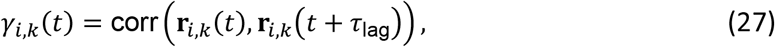

where **r**_*i,k*_(*t*) is the spatially-flattened activity vector at time *t*. We used *τ*_lag_ = 10*τ* for the model, and *τ*_lag_ = 0.6s for the experimental data). This provides a measure of temporal stability across trials.

To obtain a single stability value per trial, we computed the correlation *β*_*i,k*_ between two specific time points, *t*_1_ and *t*_2_, chosen to represent the response evolution,

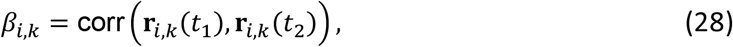

where *t*_1_ and *t*_2_ are fixed time points (model: *t*_1_ = 10*τ, t*_2_ = 40*τ* steps; experiment: *t*_1_ = 1 s, *t*_2_ = 3 s post-onset). The time points *t*_1_ and *t*_2_ were selected to capture variations in the sustained response and to correlate stability with stimulus alignment (Fig. 4F, K). This value was then averaged across all *N*_trials_ to yield a stimulus-specific intra-trial stability score, *β*_*i*_, as defined by:

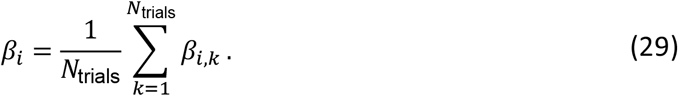

This provides a single stability metric for each stimulus pattern.

#### Similarity to stimulus (temporal fidelity)

As a second measure of intra-trial dynamics, we assessed the temporal fidelity, or how closely the spatial pattern at each time point matched the stimulus pattern (see Figure 3D, H). This time-varying similarity *κ*_*i,k*_(*t*) for stimulus *i* and trial *k* was defined as:

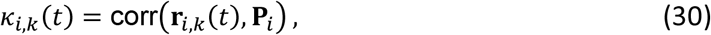

where **r**_*i,k*_(*t*) is the spatially-flattened activity vector at time *t*, and **P**_*i*_ is the (static) spatially-flattened binarized stimulus pattern. This provides a continuous measure of response fidelity over time. To obtain a single summary value for comparison, this measure was averaged across trials at a specific representative time point, *t*_spec_ so that

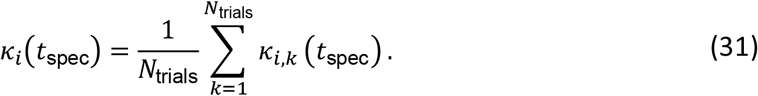

This score was evaluated at a representative steady-state time point *t*_spec_ (40*τ* for the model, 3 s post-onset for the experiment) to compare fidelity across stimulus patterns.

### Stimulus alignment with spontaneous activity

To quantify the alignment of stimulus patterns with spontaneous activity (e.g Figure 4C), we computed a measure analogous to the alignment with the connectivity modes above (Eq. 26), following the measure defined in (*29*). First, the fraction of spontaneous variance explained by each stimulus pattern **P**_*i*_ was calculated as

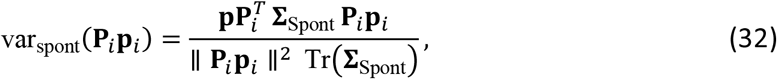

where **∑**_Spont_ is the covariance matrix of spontaneous activity, Tr(⋅) its trace, and ∥ **P**_*i*_ ∥^2^ is the squared L2-norm (equivalent to the number of stimulated pixels for a binarized pattern). This can be equivalently expressed as 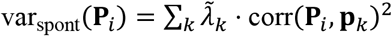, where 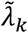 and **p**_*k*_ are normalized eigenvalues and eigenvectors of the spontaneous covariance matrix **∑**_Spont_ (i.e. explained variance ratio and principal components), replacing singular values and input singular vectors from Eq. (24).

To account for the binarized nature of the input patterns, we normalized this variance by the maximum variance a binarized pattern aligned to the first spontaneous principal component (**p**_1_) could explain. This normalization constant *c*_Norm_ was defined as

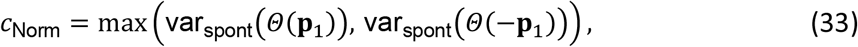

where the binarization Θ is applied to **p**_1_ and its sign-flipped version −**p**_1_. For the computational model, *c*_Norm_ was calculated separately for each network realization. For the experimental analysis, to account for large inter-animal variability in the variance explained by the first PC, we used a single normalization factor obtained by averaging Eq. (33) across all animals. The final normalized alignment score for each stimulus pattern was then

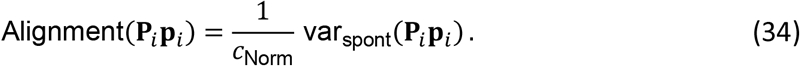

### Quantification and statistical analysis

We used re-sampling methods for hypothesis testing and statistical analysis in the computational modelling results (eg. Fig. 2B-D). Significance was assessed by comparing the average value of a given statistic between the real data and a suitable control distribution. The p-value was computed as the fraction of values in the control distribution that were more extreme than the difference observed in the real data.

#### Paired permutation test

To test whether a group of animals showed a significant change across conditions (for instance, to establish whether the difference in trial-to-trial correlation in Fig. 2D is significant), we employed a paired permutation test to generate a control distribution. We randomly flipped the data points for a given animal between conditions and recomputed the difference between the group averages. Repeating this procedure *N* = 1000 times resulted in a control distribution, which we compared with the real difference.

#### Wilcoxon Signed-Rank test

For paired comparisons where a full permutation distribution was not generated (for statistical analyses in the experimental data), we assessed significance using the two-sided Wilcoxon Signed-Rank test.

Center and spread values are reported as mean and standard error of the mean (SEM) within a given condition, unless otherwise noted. Statistical analyses were performed in Python, and significance was defined as *p* <0.05 and indicated with the symbol *.

## Supplemental figures

**Figure S1.**
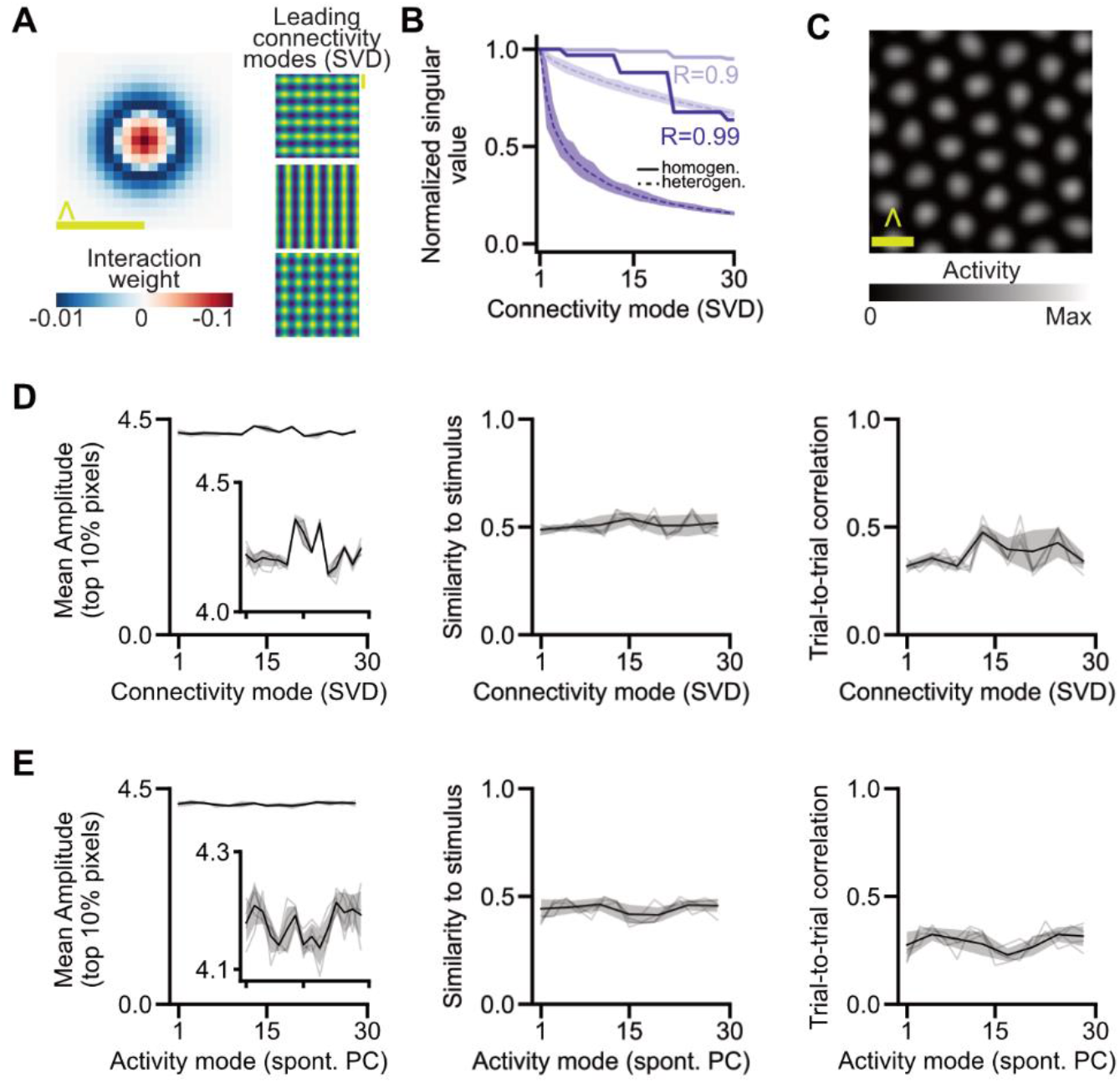
Additional analyses on selective amplification effects in nonlinear, homogeneous networks. (related to Fig 1, Fig. 4) **(A)** Recurrent network interactions with lateral excitation and long-range inhibition (LE/LI) interaction profile (*left*), and corresponding first three connectivity modes (calculated via SVD) (*right*). For a symmetric network (as here with a homogeneous connectivity profile), input and output connectivity mode patterns are identical. Yellow bar indicates intrinsic wavelength (⋀). **(B)** Associated singular values for the first thirty modes for two different recurrent strengths (see Figure 1D). **(C)** Example spontaneous activity pattern in such a homogeneous recurrent network. **(D)** Compare Fig 1 F: Peak response amplitude (*left*), similarity to stimulus (*middle*), and trial-to-trial correlation (*right*) show little change with connectivity mode order and show no clear relationship to associated singular values. Individual lines: N=3 independent networks. Thick line and shaded region show mean and standard deviation. As we simulated a finite-size system with finite integration time, defects in the optimal theoretical solution (hexagonal patterns) for that input regime remain, and we observe slight variations between the first 30 connectivity modes. **(E)** As D, but depending on activity modes – spontaneous principal components.

**Figure S2.**
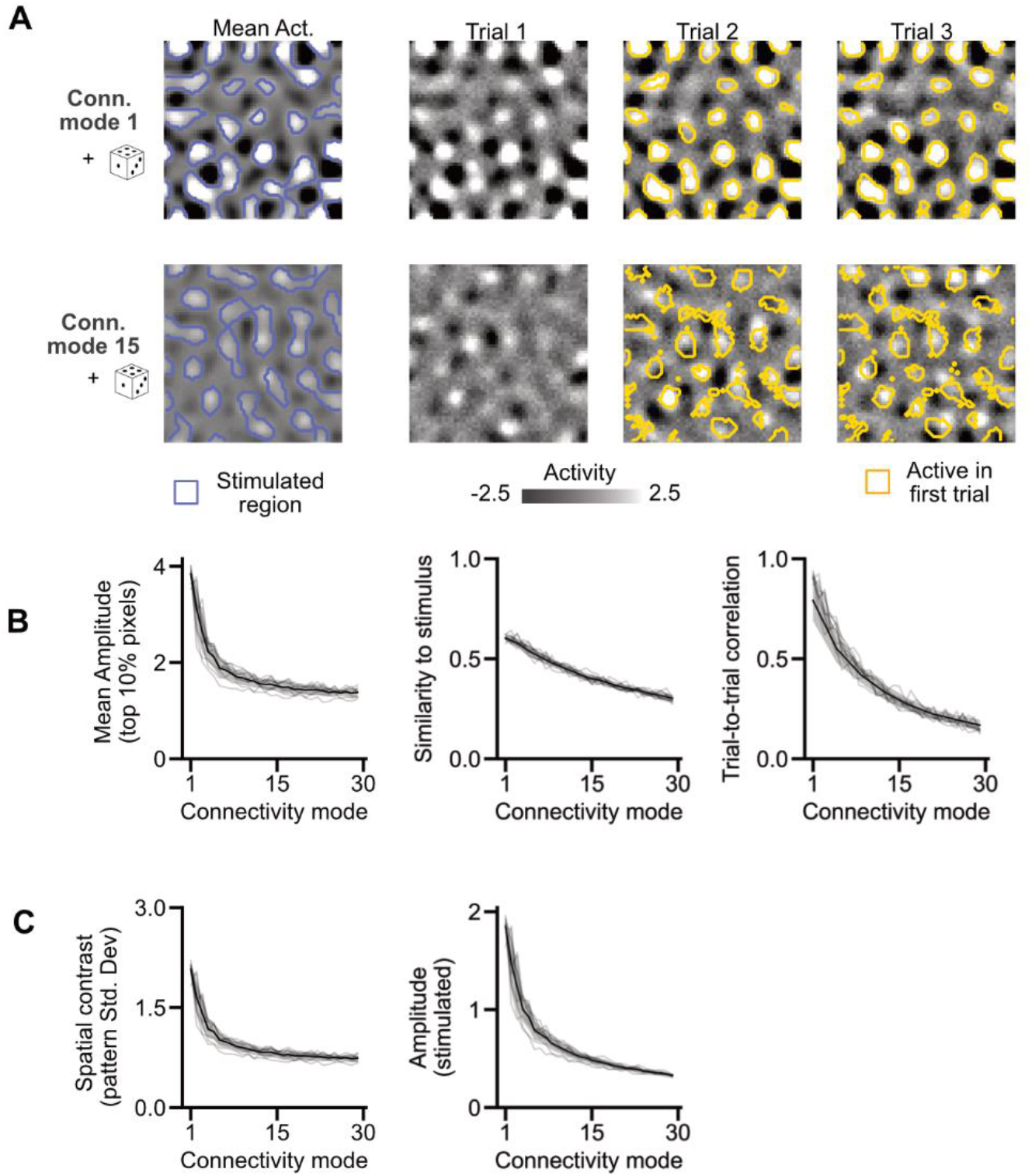
Stimulus patterns close to dominant connectivity modes drive reliable network responses *in silico* in a linear network. (related to Fig 1). Recurrent network with lateral excitation and long-range inhibition (LE/LI) interaction profile with a mild degree of additional heterogeneity, solved for steady state in a linear regime. **(A)** (*Left*) Driving network with patterns based on connectivity modes and additional noise. (*Middle*) Mean evoked network response across trials (n=40 trials), contours mark stimulated region. (*Right*) Example network activity patterns, contours mark active region in first trial. **(B)** Stimulus patterns based on dominating connectivity (high associated singular values) modes evoke responses with greater peak amplitude (*left*), similarity to stimulus (*middle*), and trial-to-trial correlation (*right*) compared non-dominant connectivity modes with weaker associated singular values. Individual lines: N=6 independent networks. Thick line and shaded region show mean and standard deviation. **(C)** Same as (B), but for spatial contrast (*left*) and mean ΔF/F amplitude within stimulated pixels (*right*).

**Figure S3.**
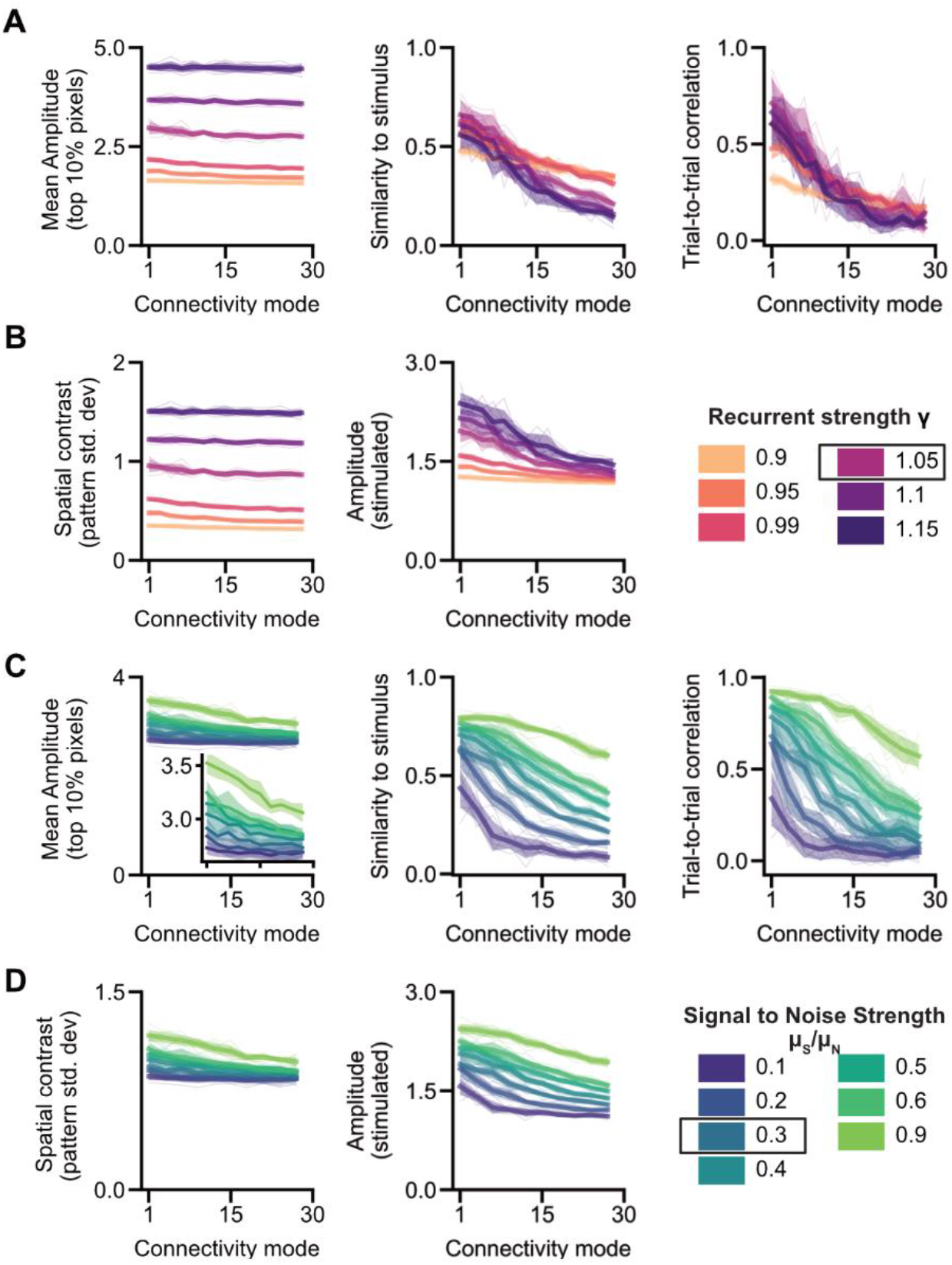
Additional analyses on selective amplification effects in nonlinear networks. (related to Fig 1) **(A-B)** Variations in recurrent strength. Value chosen for Fig. 1 indicated with black box (1.05). **(A)** As Fig 1 F: Stimulus patterns based on dominating connectivity (high associated singular values) modes evoke responses with greater amplitude (*left*), similarity to stimulus (*middle*), and trial-to-trial correlation (*right*) compared non-dominant connectivity modes with weaker associated singular values. Individual lines: N=6 independent networks. Thick line and shaded region show mean and standard deviation. Colors represent six different levels of recurrent interaction strength in a network with rectifying nonlinearity (see right hand side legend). **(B)** Additional analysis on amplitude effects (as in A), quantified with methods measuring spatial contrast (*left*), and amplitude within stimulated regions (*middle*) (see Methods). **(C-D** Variations in signal to Noise strength. Value chosen for Fig. 1 indicated with black box (0.3). Panels as A,B.

**Figure S4.**
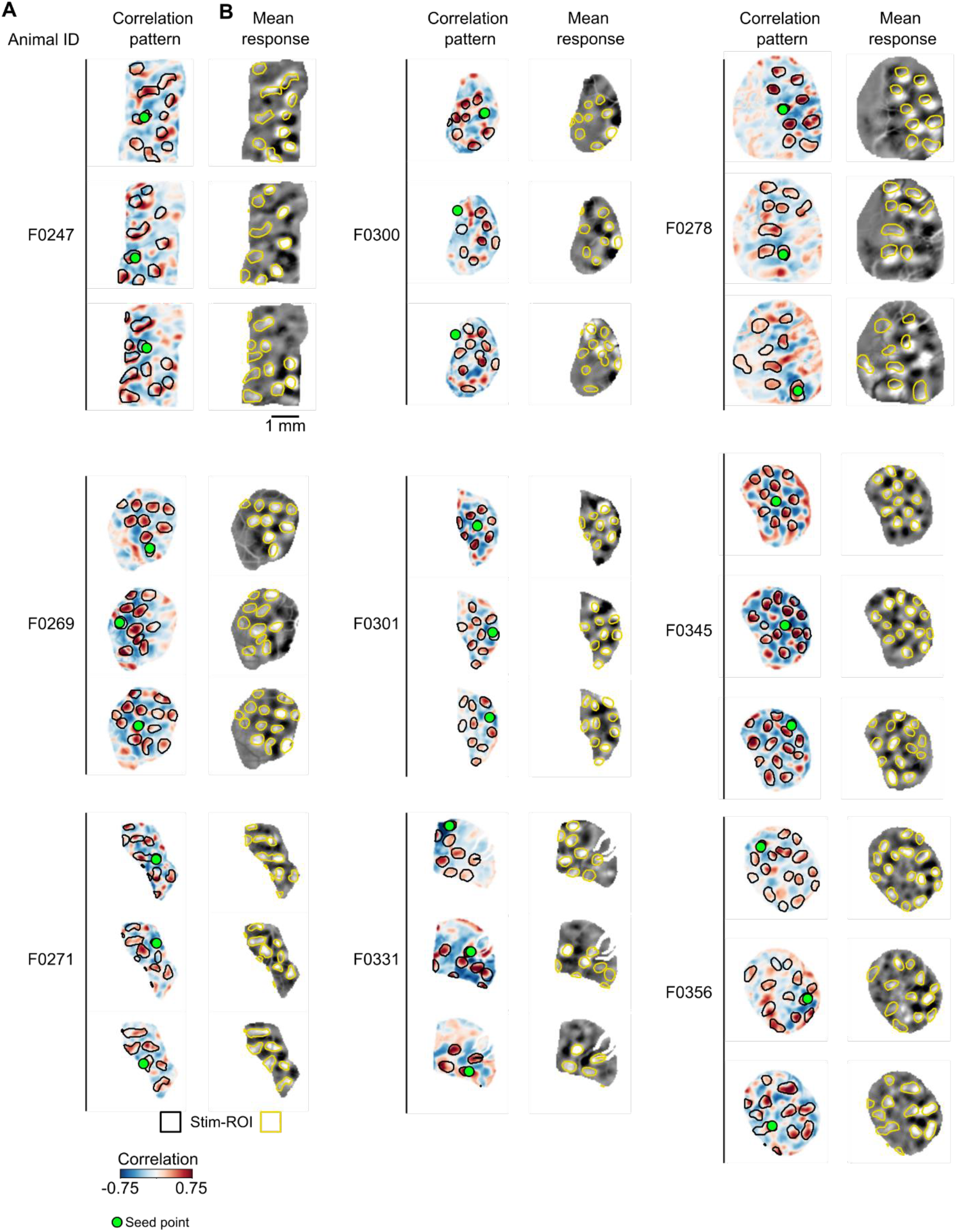
Opto-stimuli based on spontaneous correlation patterns evoke stimulus specific responses. (related to Fig. 2F,G) (**A**) For all animals, 3 different optogenetic stimuli based on endogenous activity were designed by doing online analysis of spontaneous correlation patterns conducted on the day of the experiment. Shown here are the stimulus patterns (Black contours) plotted on top of spontaneous correlation patterns from example seed points (green dots) after full post-experiment processing of the data, showing that patterns drawn on the day of the experiment are largely maintained. (**B**) The mean opto-evoked response to a given stimulus pattern (Yellow contours) for all animals, showing stimulus specific responses. For all animals, FOVs are constrained to only opto-responsive area. All FOVs scaled to the same scale bar, 1 mm.

**Figure S5.**
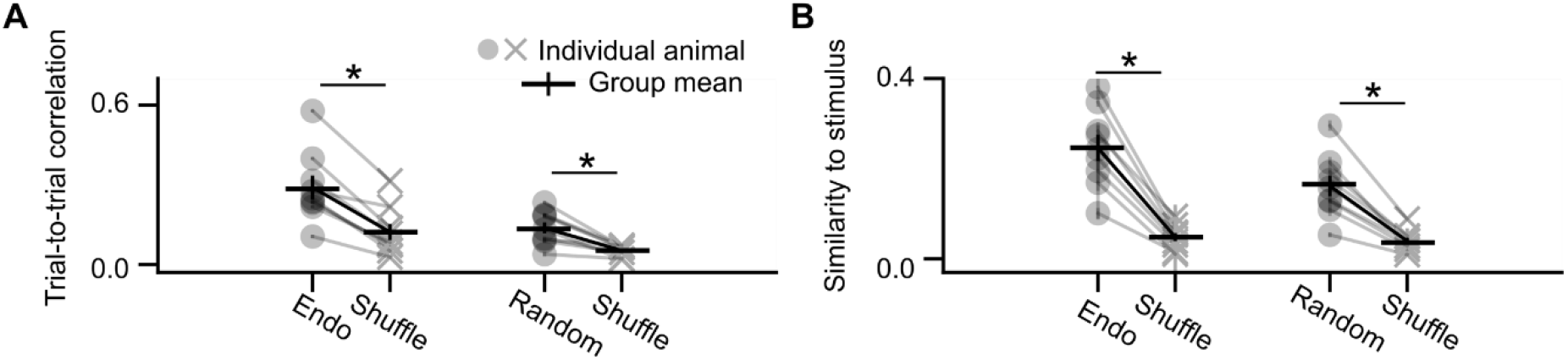
Both endogenous and random stimuli evoke reliable and specific responses. (related to Fig. 2L,M) (**A**) Endo and random stimuli have significantly greater trial-to-trial correlations than trial-shuffled controls (average across N=500 random permutations). Trial to trial correlation calculated on a per stimulus pattern basis (n=40 trials) and then averaged (+/− SEM) across stimuli (n=3) for each animal (circles, crosses). (* indicates p<0.01, WRS test). (**B**) Same as A, but for similarity to respective stimulus patterns. (* indicates p<0.01, WRS test)

**Figure S6.**
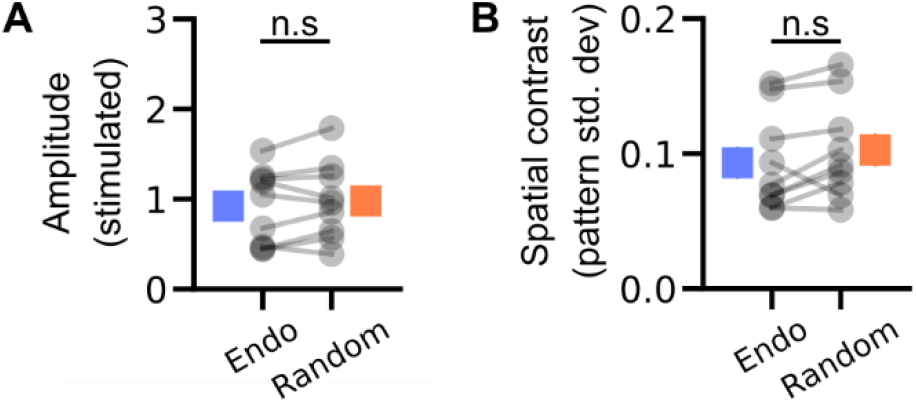
ΔF/F amplitude of endogenous and random pattern evoked responses *in vivo* (related to Figure 2). (**A**) Average ΔF/F response within stimulated pixels evoked by endogenous (blue) and random (orange) stimuli. (**B**) Average spatial contrast across FOV on a per trial basis, suggesting similar variance in the peaks and troughs of columnar activity for endogenous and random stimuli. N=9 animals, n.s. indicates p values >0.05, Wilcoxon signed rank test.

**Figure S7.**
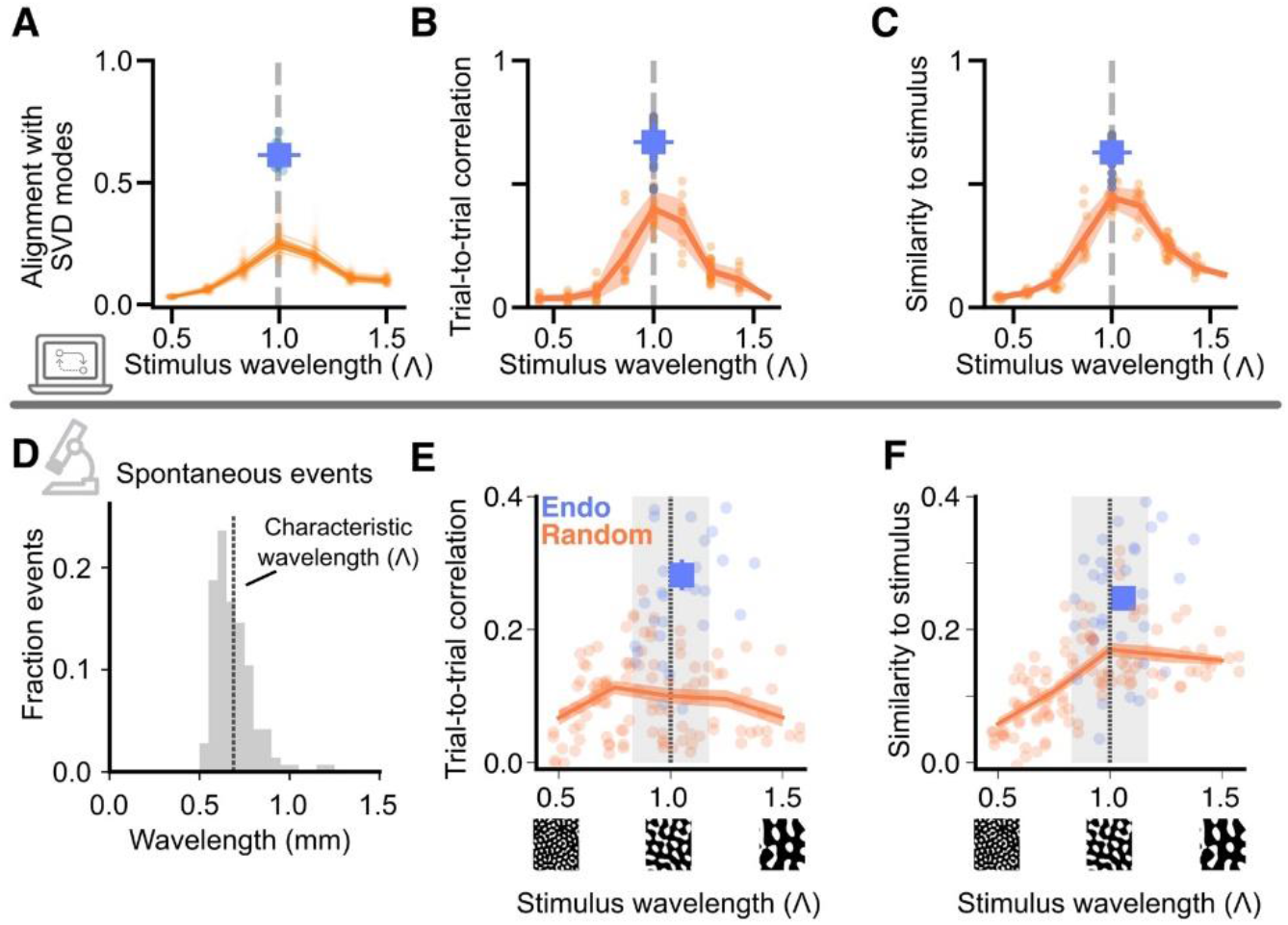
Selective amplification of endogenous modes beyond wavelength amplification (related to Fig 2). (**A**) Alignment with SVD modes for endogenous derived stimuli (blue, dots: 10 independent network simulations, errorbar: mean +/− SEM) and random patterns of different wavelengths (thin lines: 10 independent network simulations, thick line and shaded region: mean SEM). The characteristic wavelength is determined from network connectivity and spontaneous activity. (**B-C**) Computational model simulation: Average trial-to-trial correlation (**B**) and similarity to stimulus (**C**) as a function of input wavelength for endogenous (blue) and random (orange) stimuli of varying wavelength, (n=10 stimuli for 15 different center wavelengths across 10 independent network simulations). Blue square indicates mean (+/− SEM) for all endogenous inputs. Orange line indicates binned mean (+/− SEM, bin size=0.25 Λ) for all random inputs. (**D**) Example distribution of the spatial wavelength of spontaneous events from a single animal. Average spontaneous event wavelength is used to estimate the network’s characteristic wavelength (Λ). (**E**-**F**) Same as (B-C), but for *in vivo* data. n=140 patterns, pooled across 9 animals. Grey lines and bars show the mean (+/− Std. Dev) wavelength of spontaneous events across animals. Stimulus wavelength is normalized to each animal’s characteristic wavelength. (+/− SEM, bin size=0.25 Λ)

## Notes

### Competing Interest Statement

The authors have declared no competing interest.

## References

1. L. Cossell, M. F. Iacaruso, D. R. Muir, R. Houlton, E. N. Sader, H. Ko, S. B. Hofer, T. D. Mrsic-Flogel, Functional organization of excitatory synaptic strength in primary visual cortex. Nature 518, 399–403 (2015).

2. H. Ko, L. Cossell, C. Baragli, J. Antolik, C. Clopath, S. B. Hofer, T. D. Mrsic-Flogel, The emergence of functional microcircuits in visual cortex. Nature 496, 96–100 (2013).

3. I. A. Oldenburg, W. D. Hendricks, G. Handy, K. Shamardani, H. A. Bounds, B. Doiron, H. Adesnik, The logic of recurrent circuits in the primary visual cortex. Nat Neurosci 27, 137–147 (2024).

4. J. F. O’Rawe, Z. Zhou, A. J. Li, P. K. LaFosse, H. C. Goldbach, M. H. Histed, Excitation creates a distributed pattern of cortical suppression due to varied recurrent input. Neuron 111, 4086–4101.e5 (2023).

5. S. Peron, R. Pancholi, B. Voelcker, J. D. Wittenbach, H.F. Ólafsdóttir, J. Freeman, K. Svoboda, Recurrent interactions in local cortical circuits. Nature 579, 256–259 (2020).

6. I. K. Christie, P. Miller, S. D. Van Hooser, Cortical amplification models of experience-dependent development of selective columns and response sparsification. Journal of Neurophysiology 118, 874–893 (2017).

7. R. J. Douglas, C. Koch, M. Mahowald, K. A. C. Martin, H. H. Suarez, Recurrent Excitation in Neocortical Circuits. Science 269, 981–985 (1995).

8. K. D. Miller, Canonical computations of cerebral cortex. Current Opinion in Neurobiology 37, 75–84 (2016).

9. B. K. Murphy, K. D. Miller, Balanced Amplification: A New Mechanism of Selective Amplification of Neural Activity Patterns. Neuron 61, 635–648 (2009).

10. F. S. Chance, S. B. Nelson, L. F. Abbott, Complex cells as cortically amplified simple cells. Nat Neurosci 2, 277–282 (1999).

11. D. A. Aponte, G. Handy, A. M. Kline, H. Tsukano, B. Doiron, H. K. Kato, Recurrent network dynamics shape direction selectivity in primary auditory cortex. Nat Commun 12, 314 (2021).

12. Y. H. Liu, A. Baratin, J. Cornford, S. Mihalas, E. Shea-Brown, G. Lajoie, How connectivity structure shapes rich and lazy learning in neural circuits. arXiv 2310.08513 [Preprint] (2024). http://arxiv.org/abs/2310.08513.

13. F. Mastrogiuseppe, S. Ostojic, Linking Connectivity, Dynamics, and Computations in Low-Rank Recurrent Neural Networks. Neuron 99, 609–623.e29 (2018).

14. C. E. Deveau, Z. Zhou, P. K. LaFosse, Y. Deng, S. Mirbagheri, N. Steinmetz, M. H. Histed, Recurrent cortical networks encode natural sensory statistics via sequence filtering. Neuron, doi: 10.1016/j.neuron.2025.12.024. (2026).

15. J. H. Marshel, Y. S. Kim, T. A. Machado, S. Quirin, B. Benson, J. Kadmon, C. Raja, A. Chibukhchyan, C. Ramakrishnan, M. Inoue, J. C. Shane, D. J. McKnight, S. Yoshizawa, H. E. Kato, S. Ganguli, K. Deisseroth, Cortical layer–specific critical dynamics triggering perception. Science 365, eaaw5202 (2019).

16. T. Bonhoeffer, A. Grinvald, Iso-orientation domains in cat visual cortex are arranged in pinwheel-like patterns. Nature 353, 429–431 (1991).

17. B. Chapman, M. P. Stryker, T. Bonhoeffer, Development of Orientation Preference Maps in Ferret Primary Visual Cortex. The Journal of Neuroscience 16, 6443–6453 (1996).

18. D. H. Hubel, T. N. Wiesel, Receptive Fields and Functional Architecture of Monkey Striate Cortex. Journal of Physiology 195, 215–243 (1968).

19. V. Mountcastle, The columnar organization of the neocortex. Brain 120, 701–722 (1997).

20. A. A. Lempel, S. Trägenap, C. Tepohl, M. Kaschube, D. Fitzpatrick, Development of coherent cortical responses reflects increased discriminability of feedforward inputs and their alignment with recurrent circuits. Neuron, S0896627325005999 (2025).

21. T. Rózsa, R. Cagnol, J. Antolík, Iso-orientation bias of layer 2/3 connections: the unifying mechanism of spontaneous, visually and optogenetically driven V1 dynamics. [Preprint] (2025). 10.1101/2024.11.19.624284.

22. N. J. Powell, B. Hein, D. Kong, J. Elpelt, H. N. Mulholland, M. Kaschube, G. B. Smith, Common modular architecture across diverse cortical areas in early development. Proceedings of the National Academy of Sciences 121, 1–10 (2024).

23. N. J. Powell, B. Hein, D. Kong, J. Elpelt, H. N. Mulholland, R. A. Holland, M. Kaschube, G. B. Smith, Developmental maturation of millimeter-scale functional networks across brain areas. Cerebral Cortex 35, bhaf007 (2025).

24. H. N. Mulholland, H. Jayakumar, D. M. Farinella, G. B. Smith, All-optical interrogation of millimeter-scale networks and application to developing ferret cortex. Journal of Neuroscience Methods, doi: 10.1016/j.jneumeth.2023.110051 (2023).

25. H. N. Mulholland, M. Kaschube, G. B. Smith, Self-organization of modular activity in immature cortical networks. Nat Commun 15, 4145 (2024).

26. G. B. Smith, B. Hein, D. E. Whitney, D. Fitzpatrick, M. Kaschube, Distributed network interactions and their emergence in developing neocortex. Nature Neuroscience 21, 1600–1608 (2018).

27. L. F. Abbott, Decoding neuronal firing and modelling neural networks. Quart. Rev. Biophys. 27, 291–331 (1994).

28. S. H. Seung, “Amplification, Attenuation, and Integration” in The Handbook of Brain Theory and Neural Networks / Edited by Michael A. Arbib (MIT Press, ed. 2, 2003; https://direct.mit.edu/books/edited-volume/4358/The-Handbook-of-Brain-Theory-and-Neural-Networks), pp. 94–97.

29. S. Trägenap, D. E. Whitney, D. Fitzpatrick, M. Kaschube, The developmental emergence of reliable cortical representations. Nat Neurosci 28, 394–405 (2025).

30. M. Kaschube, M. Schnabel, S. Lowel, D. M. Coppola, L. E. White, F. Wolf, Universality in the Evolution of Orientation Columns in the Visual Cortex. Science 330, 1113–1116 (2010).

31. C. von der Malsburg, Self-Organization of Orientation Sensitive Cells in the Striate Cortex. Biological Cybernetics 14, 85–100 (1973).

32. G. B. Ermentrout, J. D. Cowan, A Mathematical Theory of Visual Hallucination Patterns. Biol. Cybernetics 34, 137–150 (1979).

33. U. A. Ernst, K. R. Pawelzik, C. Sahar-Pikielny, M. V. Tsodyks, Intracortical origin of visual maps. Nat Neurosci 4, 431–436 (2001).

34. W. H. Bosking, Y. Zhang, B. Schofield, D. Fitzpatrick, Orientation selectivity and the arrangement of horizontal connections in tree shrew striate cortex. The Journal of Neuroscience 17, 2112–27 (1997).

35. A. Angelucci, J. B. Levitt, E. J. S. Walton, J.-M. Hupé, J. Bullier, J. S. Lund, Circuits for Local and Global Signal Integration in Primary Visual Cortex. J. Neurosci. 22, 8633–8646 (2002).

36. S. Panzeri, J. H. Macke, J. Gross, C. Kayser, Neural population coding: combining insights from microscopic and mass signals. Trends in Cognitive Sciences 19, 162–172 (2015).

37. K. E. Schmidt, S. Löwel, “Long-Range Intrinsic Connections in Cat Primary Visual Cortex” in The Cat Primary Visual Cortex (Academic Press, 2002; 10.1016/B978-012552104-8/50011-0), pp. 387–426.

38. S. Kira, H. Safaai, A. S. Morcos, S. Panzeri, C. D. Harvey, A distributed and efficient population code of mixed selectivity neurons for flexible navigation decisions. Nat Commun 14, 2121 (2023).

39. A. Kohn, R. Coen-Cagli, I. Kanitscheider, A. Pouget, Correlations and Neuronal Population Information. Annu. Rev. Neurosci. 39, 237–256 (2016).

40. T. Kenet, D. Bibitchkov, M. Tsodyks, A. Grinvald, A. Arieli, Spontaneously emerging cortical representations of visual attributes. Nature 425, 954–956 (2003).

41. K. Korvasová, F. Grani, M. Voldřich, R. López Peco, D. Berling, M. Val Calvo, A. Rodil Doblado, T. Rózsa, C. Soto Sánchez, X. Chen E. Fernandez, J. Antolík, Contributed Talks I: Recruiting native visual representations in visual cortex for electrode array based vision restoration. Journal of Vision 25 (2025).

42. A. Nejatbakhsh, F. Fumarola, S. Esteki, T. Toyoizumi, R. Kiani, L. Mazzucato, Predicting the effect of micro-stimulation on macaque prefrontal activity based on spontaneous circuit dynamics. Phys. Rev. Research 5, 043211 (2023).

43. J. A. Gallego, M. G. Perich, L. E. Miller, S. A. Solla, Neural Manifolds for the Control of Movement. Neuron 94, 978–984 (2017).

44. M. D. Golub, P. T. Sadtler, E. R. Oby, K. M. Quick, S. I. Ryu, E. C. Tyler-Kabara, A. P. Batista, S. M. Chase, B. M. Yu, Learning by neural reassociation. Nat Neurosci 21, 607–616 (2018).

45. E. R. Oby, M. D. Golub, J. A. Hennig, A. D. Degenhart, E. C. Tyler-Kabara, B. M. Yu, S. M. Chase, A. P. Batista, New neural activity patterns emerge with long-term learning. Proc. Natl. Acad. Sci. U.S.A. 116, 15210–15215 (2019).

46. P. T. Sadtler, K. M. Quick, M. D. Golub, S. M. Chase, S. I. Ryu, E. C. Tyler-Kabara, B. M. Yu, A. P. Batista, Neural constraints on learning. Nature 512, 423–426 (2014).

47. A. Roy, S. Wang, B. Meschede-Krasa, J. Breffle, S. D. Van Hooser, An early phase of instructive plasticity before the typical onset of sensory experience. Nat Commun 11, 11 (2020).

48. J. Antolík, R. Cagnol, T. Rózsa, C. Monier, Y. Frégnac, A. P. Davison, A comprehensive data-driven model of cat primary visual cortex. PLoS Comput Biol 20, e1012342 (2024).

49. D. Ferster, K. D. Miller, Neural Mechanisms of Orientation Selectivity in the Visual Cortex. Annu. Rev. Neurosci. 23, 441–471 (2000).

50. E. Margalit, H. Lee, D. Finzi, J. J. DiCarlo, K. Grill-Spector, D. L. K. Yamins, A unifying framework for functional organization in early and higher ventral visual cortex. Neuron 112, 2435–2451.e7 (2024).

51. G. B. Smith, D. Fitzpatrick, Viral Injections and Cranial Window Implantation for In Vivo Two-Photon Imaging. Methods Mol Biol 1474, 171–185 (2016).

52. H. N. Mulholland, B. Hein, M. Kaschube, G. B. Smith, Tightly coupled inhibitory and excitatory functional networks in the developing primary visual cortex. eLife 10 (2021).

53. H. R. Wilson, J. D. Cowan, Excitatory and Inhibitory Interactions in Localized Populations of Model Neurons. Biophysical Journal 12, 1–24 (1972).

54. H. R. Wilson, J. D. Cowan, A mathematical theory of the functional dynamics of cortical and thalamic nervous tissue. Kybernetik 13, 55–80 (1973).

